# Categorical Biases in Human Occipitoparietal Cortex

**DOI:** 10.1101/170845

**Authors:** Edward F. Ester, Thomas C. Sprague, John T. Serences

## Abstract

Categorization allows organisms to generalize existing knowledge to novel stimuli and to discriminate between physically similar yet conceptually different stimuli. Humans, nonhuman primates, and rodents can readily learn arbitrary categories defined by low-level visual features, and learning distorts perceptual sensitivity for category-defining features such that differences between physically similar yet categorically distinct exemplars are enhanced while differences between equally similar but categorically identical stimuli are reduced. We report a basis for these distortions in human occipitoparietal cortex. In three experiments, we used an inverted encoding model to recover population-level representations of stimuli from multivoxel and multi-electrode patterns of human brain activity while human participants (both sexes) classified continuous stimulus sets into discrete groups. In each experiment, reconstructed representations of to-be-categorized stimuli were systematically biased towards the center of the appropriate category. These biases were largest for exemplars near a category boundary, predicted participants’ overt category judgments, emerged shortly after stimulus onset, and could not be explained by mechanisms of response selection or motor preparation. Collectively, our findings suggest that category learning can influence processing at the earliest stages of cortical visual processing.

**Significance Statement:** Category learning enhances perceptual sensitivity for physically similar yet categorically different stimuli. We report a possible mechanism for these distortions in human occipitoparietal cortex.. In three experiments, we used an inverted encoding model to recover population-level representations of stimuli from multivariate patterns in occipitoparietal cortex while participants categorized sets of continuous stimuli into discrete groups. The recovered representations were systematically biased by category membership, with larger biases for exemplars adjacent to a category boundary. These results suggest that mechanisms of categorization shape information processing at the earliest stages of the visual system.

---

Categorization refers to the process of mapping continuous sensory inputs onto discrete and behaviorally relevant concepts. It is a cornerstone of flexible behavior that allows organisms to generalize existing knowledge to novel stimuli and to discriminate between physically similar yet conceptually different stimuli. Many real-world categories are defined by a combination of low-level visual properties such as hue, luminance, spatial frequency, and orientation. For example, a forager might be tasked with determining whether a food source is edible based on subtle variations in color, shape, size, and texture. Humans and other animals can readily learn arbitrary novel categories defined by low-level visual properties (Goldstone, 1998; Ashby & Maddox, 2005), and such learning “distorts” perceptual sensitivity for category-defining features such that discrimination performance for physically similar yet categorically different stimuli is increased (i.e., acquired distinctiveness; Goldstone, 1995; Newell & Bulthoff, 2002) and discrimination performance for stimuli from the same category reduced (i.e., acquired similarity; Livingston et al., 1998).

Invasive electrophysiological studies suggest that single-unit responses in early visual areas index the physical properties of a stimulus but not its category membership, while single-unit responses in later areas index the category membership of a stimulus regardless of its physical properties (e.g., Sigala & Logothetis, 2002; Freedman et al., 2001; Freedman & Assad, 2006). These results have been taken as evidence that category-selective responses are a *de novo* property of higher-order visual areas. However, perceptual distortions following category learning could also reflect subtle changes in how to-be-categorized information is represented by sensory neural populations (Folstein et al., 2012; Davis & Poldrack, 2013). Here we provide a test of this possibility. In three experiments, we trained human participants (both sexes) to classify sets of continuous stimuli into discrete groups. Next, next, we applied multivariate models to noninvasive measurements of human brain activity (fMRI and EEG) from visual and parietal cortical areas while participants categorized the same stimulus sets. This allowed us to recover, visualize, and quantify stimulus-specific representations of to-be-categorized exemplars. In Experiment 1 (fMRI), we show that reconstructed representations of to-be-categorized orientations in visual areas V1-V3 are systematically biased towards the center of the category to which they belong. These biases were correlated with trial-by-trial variability in overt category judgments and were largest for orientations adjacent to the category boundary where they would be most beneficial for discrimination performance. In Experiment 2, we utilized EEG to generate time-resolved representations of to-be-categorized orientations and show that categorical biases manifest shortly after stimulus onset (≤ 300 ms). In Experiment 3, we used EEG and a delayed match-to-category task to show that categorical biases observed in Experiments 1 and 2 cannot be explained by response biases or motor preparation. Collectively, our findings suggest that mechanisms of categorization can shape information processing at the earliest stages of the visual system.

## Methods

### General Overview

#### Participants

A total of 44 human volunteers (both sexes) participated in this study. Eight participants completed Experiment 1 (fMRI), 28 participants completed Experiment 2 (EEG), and eight participants completed Experiment 3 (EEG). Experiments 1 and 2 were performed at the University of California, San Diego, while Experiment 3 was performed at Florida Atlantic University. Participants were recruited from the student body at each university. All study procedures were approved by local institutional review boards, and all participants gave both written and oral informed consent. Participants self-reported normal or corrected-to-normal visual acuity and were remunerated with cash incentives ($20/h for fMRI and $15/h for EEG).

#### Stimulus Displays

Stimulus displays were generated in MATLAB and rendered using Psychophysics Toolbox software extensions (Kleiner et al., 2017). During Experiment 1 (fMRI), displays were projected onto a 110 cm-wide screen placed at the base of the MRI table, and participants viewed displays via a mirror attached to the MR head coil from a distance of 370 cm. During Experiments 2 and 3, displays were projected onto a 19-inch CRT monitor cycling at 120Hz (Experiment 2) or 85Hz (Experiment 3). Participants were seated approximately 65 cm from the display (head position was not constrained).

### Experiment 1 - fMRI

#### Participants

Eight neurologically intact human volunteers (AA, AB, AC, AD, AE, AF, AG, and AH; six females) completed Experiment 1. Each participant completed a single one-hour behavioral training session approximately 24-72 hours prior to scanning. Seven participants (AA, AB, AC, AD, AE, AF, AG) completed two 2-hour experimental scan sessions; an eighth participant (AH) completed a single 2-hour experimental scan session. Participants AA, AB, AC, AD, AE, AF, and AH also completed a single 2-hour retinotopic mapping scan session. Data from this session were used to identify visual field borders in early visual cortical areas V1-hV4/V3A and subregions of posterior intraparietal sulcus (IPS0-3; see *Retinotopic Mapping*, below)

#### Behavioral Tasks

In separate runs (where “run” refers to a continuous block of 30 trials lasting 280 seconds) participants performed either an orientation mapping task or a category discrimination task. Trials in each task lasted 3 seconds, and consecutive trials were separated by a 5 or 7 s inter-trial-interval (pseudorandomly chosen on each trial). During the orientation mapping task, participants attended a stream of letters presented at fixation (subtending 1.0° × 1.0° from a viewing distance of 370 cm) while ignoring a task-irrelevant phase-reversing (15 Hz) square-wave grating (0.8 cycles/deg with inner and outer radii of 1.16° and 4.58°, respectively) presented in the periphery. On each trial, the grating was assigned one of 15 possible orientations (0°-168° in 12° increments). Participants were instructed to detect and report the identity of a target (“X” or “Y”) in the letter stream using an MR-compatible button box. Only one target was presented on each trial. Letters were presented at a rate of 10 Hz (50% duty cycle, i.e. 50 msec on, 50 msec off), and targets could occur during any cycle from +750 to +2250 msec after stimulus onset. During category discrimination runs, participants were shown displays containing a circular aperture (inner and outer radii of 1.16° and 4.58° from a viewing distance of 370 cm) filled with 150 iso-oriented bars (see Figure 1A). Each bar subtended 0.2° × 0.6° with a stroke width of 8 pixels (1024 × 768 display resolution). Each bar flickered at 30 Hz and was randomly replotted within the aperture at the beginning of each “up” cycle.

**Figure 1.**
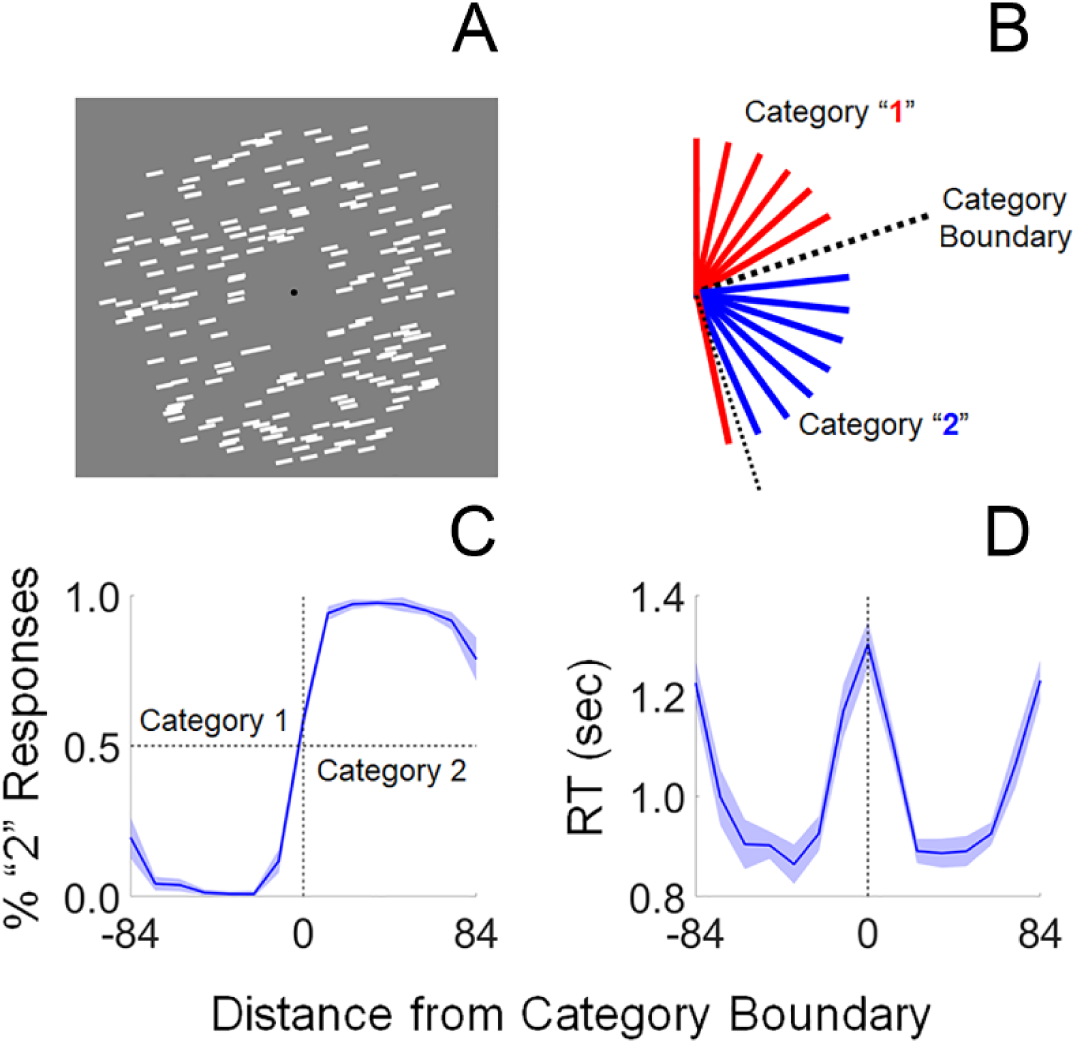
Overview of Experiment 1. (A) Participants viewed displays containing a circular aperture of iso-oriented bars. On each trial, the bars were assigned one of 15 unique orientations from 0-168°. (B) We randomly selected and designated one stimulus orientation as a category boundary (black dashed line), such that the seven orientations counterclockwise from this value were assigned to Category 1 (red lines) and the seven orientations clockwise from this value were assigned to Category 2 (blue lines). (C) After training, participants rarely miscategorized orientations. (D) Response latencies are significantly longer for oriented exemplars near the category boundary (RT = response time; shaded regions in C-D are ±1 within-participant S.E.M.).

On each trial, all bars were assigned an orientation from 0°-168° in 12° increments. Inspired by earlier work in non-human primates (Freedman & Assad, 2006), we randomly selected and designated one of these orientations as a category boundary such that the seven orientations counterclockwise to this value were assigned membership in “Category 1”, while the seven orientations clockwise to this value were assigned membership in “Category 2”. Participants were not informed that the category boundary was chosen from the set of possible stimulus orientations. Participants reported whether the orientation shown on each trial was a member of Category 1 or 2 (via an MR-compatible button box). Participants were free to respond at any point during the trial, though the stimulus was always presented for a total of 3000 ms. Each participant was familiarized and trained to criterion performance on the category discrimination task during a one-hour behavioral testing session completed one to three days prior to his or her first scan session. Written feedback (“Correct!” or “Incorrect”) was presented in the center of the display for 1.25 sec. after each trial during behavioral training and MR scanning. Across either one (N = 1) or two (N = 7) scan sessions, each participant completed 7 (N = 1), 13 (N = 1), 14 (N = 1), 15 (N = 1) or 16 (N = 4) runs of the orientation mapping and category discrimination tasks.

#### fMRI Acquisition and Preprocessing

Imaging data were acquired with a 3.0T GE MR 750 scanner located at the Center for Functional Magnetic Resonance imaging on the UCSD campus. All images were acquired with a 32 channel Nova Medical head coil (Wilmington, MA). Whole-brain echo-planar images (EPIs) were acquired in 35 3 mm slices (no gap) with an in-plane resolution of 3 × 3 mm (192 × 192 mm field-of-view, 64 × 64 mm image matrix, 90° flip angle, 2000 ms TR, 30 ms TE). During retinotopic mapping scans (see below) EPIs were acquired in 31 3mm thick oblique slices (no gap) positioned over posterior visual and parietal cortex with an in-plane resolution of 2 × 2 mm (192 × 192 mm field-of-view, 96 × 96 mm image matrix, 90° flip angle, 2250 ms TR, 30 ms TE). EPIs were coregistered to a high-resolution anatomical image collected during the same session (FSPGR T1-weighted sequence, 11 ms TR, 3.3 ms TE, 1100 ms TI, 172 slices, 18° flip angle, 1 mm^3^ resolution), unwarped (FSL software extensions), slice-time-corrected, motion-corrected, high-pass-filtered (to remove first-, second- and third-order drift), transformed to Talairach space, and normalized (z-score) on a scan-by-scan basis. Data from data from scan sessions were then co-registered to a high-resolution anatomical image (FSPGR T1-weighted sequences; parameters as described above) collected during the retinotopic mapping session.

#### Retinotopic Mapping

Retinotopically organized visual areas V1-hV4v/V3A were defined using data from a single retinotopic mapping run collected during each experimental scan session. Participants fixated a small dot at fixation while phase-reversing (8 Hz) checkerboard wedges subtending 60° of polar angle (at maximum eccentricity) were presented along the horizontal or vertical meridian (alternating with a period of 40 seconds; i.e., 20 seconds of horizontal stimulation followed by 20 seconds of vertical stimulation). To identify visual field borders, we constructed a general linear model with two boxcar regressors, one marking epochs of vertical stimulation and another marking epochs of horizontal stimulation. Each regressor was convolved with a canonical hemodynamic function (“double gamma” as implemented in BrainVoyager QX). Next, we generated a statistical parametric map marking voxels with larger responses during epochs of vertical relative to horizontal stimulation. This map was projected onto a computationally inflated representation of each participant’s cortical surface for visualization to aid in the definition of the borders of visual areas V1, V2v, V2d, V3v, V3d, hV4v, and V3A. Data from V2v and V2d were combined into a single V2 ROI, and data from V3v and V3d were combined into a single V3 ROI. ROIs were also combined across cortical hemispheres (e.g., left and right V1) as no asymmetries were observed and the stimulus was presented in the center of the visual field.

Seven participants (AA, AB, AC, AD, AE, AF, and AH) completed a separate two-hour retinotopic mapping scan; data from this session were used to identify retinotopically organized regions of inferior parietal sulcus (IPS0-3). During each task run, participants were shown displays containing a rotating wedge stimulus (period 24.75 or 36 sec) that subtended 72° of polar angle with inner and outer radii of 1.75 and 8.75°, respectively. In alternating blocks, the wedge contained a 4 Hz phase-reversing checkerboard or field of moving dots and participants were required to detect small, brief, and temporally unpredictable changes in checkerboard contrast or dot speed. Six participants completed between 8 and 14 task runs. To compute the best polar angle for each voxel in IPS we shifted the signals from counterclockwise runs by twice the estimated hemodynamic response function (HRF) delay (2 × 6.75 s = 13.5 s), removed data from the first and last full stimulus cycle, and reversed the time series so that all runs reflected clockwise rotation. We next computed the power and phase of the response at the stimulus’ period (either 1/24.75 or 1/36 Hz) and subtracted the estimated hemodynamic response function delay (6.75 seconds) to align the signal phase in each voxel with the stimulus’ location. Maps of orientation preference (computed via cross-correlation) were projected onto a computationally inflated representation of each participant’s grey-white matter boundary to aide in the identification of visual field borders separating IPS0-3. An eighth participant (AG) chose not to participate in an additional retinotopic mapping session. For this participant, we estimated visual field borders for visual areas V1-hV4/V3A. using data from the retinotopic mapping run collected during the participant’s sole experimental session. We did not attempt to define IPS regions IPS0-3 for this participant.

#### Decoding Categorical Biases in Visual Cortex

We used a linear decoder to examine whether fMRI activation patterns evoked by exemplars adjacent to the category boundary and at the center of each category were more similar during the category discrimination task relative to the orientation mapping task (i.e., acquired similarity). In the first phase of the analysis, we trained a linear support vector machine (LIBSVM implementation; Chang & Lin, 2011) to discriminate between the oriented exemplars at the center of each category (48° from the boundary) using data from the orientation mapping and category discrimination tasks. To ensure internal reliability, we implemented a “leave-one-run-out” cross validation scheme where data from all but one scanning run was used to train the classifier and data from the remaining scanning run were used for validation. This procedure was repeated until data from each scan had served as the validation set, and the results were averaged across permutations. Next, we trained a second classifier on activation patterns evoked by exemplars at the center of each category boundary and used the trained classifier to predict the category membership of exemplars adjacent to the category boundary. If category learning increases the similarity of activation patterns evoked by exemplars within the same category, then within-category decoding performance should be superior during the category discrimination task relative to the orientation mapping task.

#### Inverted Encoding Model of Orientation Selectivity

A linear inverted encoding model (IEM) was used to recover a model-based representation of stimulus orientation from multivoxel activation patterns measured in early visual areas (Brouwer & Heeger, 2011). The same general approach was used during Experiments 1 (fMRI) and 2 (EEG). Specifically, we modeled the responses of voxels (electrodes) measured during the orientation mapping task as a weighted sum of 15 orientation-selective channels, each with an idealized response function (half-wave-rectified sinusoid raised to the 14^th^ power). The maximum response of each channel was set to unit amplitude; thus units of response are arbitrary. Let B_1_ (*m* voxels or electrodes × *n_1_* trials) be the response of each voxel (electrode) during each trial of the RSVP task, let C_1_ (*k* filters × *n_1_* trials) be a matrix of hypothetical orientation filters, and let W (*m* voxels or electrodes × *k* filters) be a weight matrix describing the mapping between B_1_ and C_1_:

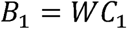

In the first phase of the analysis, we computed the weight matrix W from the voxel-wise (electrode-wise) responses in B_1_ via ordinary least-squares:

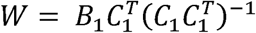

Next, we defined a test data set B_2_ (*m* voxels or electrodes × *n_2_* trials) using data from the category discrimination task. Given W and B_2_, a matrix of filter responses C_2_ (*k* filters × *n* trials) can be estimated via model inversion:

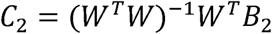

C_2_ contains the reconstructed response of each modeled orientation channel (the channel response function; CRF) on each trial of the category discrimination task. This analysis can be considered a form of model-based, directed dimensionality reduction where activity patterns are transformed from their original measurement space (fMRI voxels; EEG electrodes) into a modeled information space (orientation-selective channels). Importantly, results from this method cannot be used to infer any changes in orientation tuning – or any properties of neural responses - occurring at the single neuron level, and only assay the information content of large-scale patterns of neural activity (Sprague et al., 2018) Additionally, while it is the case that arbitrary linear transforms can be applied to the basis set, model weights, and reconstructed channel response function (Gardiner & Liu, 2019), results are uniquely defined for a given model specification (Sprague, Boynton & Serences, 2019). Trial-by-trial CRFs were multiplied by the original basis set to recover a full 180-degree function, circularly shifted to a common center (0°) and sorted by category membership so that any category bias would manifest as a clockwise shift (i.e., towards the center of Category 2).

#### Quantification of Bias in Orientation Representations

To quantify categorical biases in reconstructed model-based CRFs, these functions were fit with an exponentiated cosine function of the form:

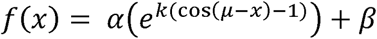

where, *x* is a vector of channel responses and α, β, *k* and µ correspond to the amplitude (i.e., signal over baseline), baseline, concentration (the inverse of bandwidth) and the center of the function, respectively. Fitting was performed using a multidimensional nonlinear minimization algorithm (Nelder-Mead).

Category biases in the estimated center of each construction (*µ*) during the category discrimination task were quantified via permutation tests. For a given visual area (e.g., V1) we randomly selected (with replacement) stimulus reconstructions from eight of eight participants. Specifically, we computed a “mean” reconstruction by randomly selecting (with replacement) and averaging reconstructions from all participants. The mean reconstruction was fit with the cosine function described above, yielding point estimates of α, β, *k*, and *µ*. This procedure was repeated 1,000 times, yielding 1,000 element distributions of parameter estimates. We then computed the proportion of permutations where a µ value less than 0 was obtained to obtain an empirical *p-*value for categorical shifts in reconstructed representations.

#### Searchlight Decoding of Category Membership

We used a roving searchlight analysis (Ester et al., 2015) to identify cortical regions beyond V1-V3 that contained category-specific information. We defined a spherical neighborhood with a radius of 8 mm around each grey matter voxel in the cortical sheet. We next extracted and averaged the normalized response of each voxel in each neighborhood over a period from 4-8 seconds after stimulus onset (this interval was chosen to account for typical hemodynamic lag of 4-6 seconds). A linear SVM (LIBSVM implementation) was used to classify stimulus category using activation patterns within each neighborhood. To classify category membership, we designated the three orientations immediately counterclockwise to the category boundary (see Figure 1) as members of Category 1 and the three orientations immediately clockwise of the boundary as members of Category 2. We then trained our classifier to discriminate between categories using data from all but one task run. The trained classifier was then used to predict category membership from activation patterns measured during the held-out task run. This procedure was repeated until each task run had been held out, and the results were averaged across permutations. Finally, we repeated the same analysis using the three Category 1 and Category 2 orientations adjacent to the second (orthogonal) category boundary (see Figure 1) and averaged the results across category boundaries.

We identified neighborhoods encoding stimulus category using a leave-one-participant-out cross validation approach (Esterman et al., 2010). Specifically, for each participant (e.g., AA) we randomly selected (with replacement) and averaged classifier performance estimates from each neighborhood from each of the remaining 7 volunteers (e.g., AB-AH). This procedure was repeated 1000 times, yielding a set of 1000 classifier performance estimates for each neighborhood. We generated a statistical parametric map (SPM) for the held-out participant that indexed neighborhoods where classifier performance was greater than chance (50%) on 97.5% of permutations (false-discovery-rate corrected for multiple comparisons across neighborhoods). Finally, we projected each participant’s SPM onto a computationally inflated representation of his or her grey-white matter boundary and used Brain Voyager’s “Create POIs from Map Clusters” function with an area threshold of 25 mm^2^ to identify ROIs supporting above-chance category classification performance. Because of differences in cortical folding patterns, some ROIs could not be unambiguously identified in all 8 participants. Therefore, across participants, we retained all ROIs that were shared by at least 7 out of 8 participants. Finally, we extracted multivoxel activation patterns from each ROI and computed model-based reconstructions of channel response functions during the RSVP and category tasks using a leave-one-run-out cross-validation approach. Specifically, we used data from all but one task run to estimate a set of orientation weights for each voxel in each ROI. We then used these weights and activation patterns measured during the held-out task run to estimate a channel response function, which contains a representation of stimulus orientation. This procedure was repeated until each task run had been held out, and the results were averaged across permutations. Note that each participant’s ROIs were defined using data from the remaining 7 participants. This ensured that participant-level reconstructions were statistically independent of the searchlight method used to define ROIs encoding category information.

#### Within-participant Error Bars

We report estimates of within-participant variability (e.g., ±1 S.E.M.) throughout the paper. These estimates discard subject variance (e.g., overall differences in BOLD response amplitude) and instead reflect variance related to the subject by condition(s) interaction term(s) (i.e., variability in estimated channel responses). We used the approach described by Cousineau (2005): raw data (e.g., channel response estimates) were de-meaned on a participant by participant basis, and the grand mean across participants was added to each participant’s zero-centered data. The grand mean-centered data were then used to compute estimates of standard error.

### Experiment 2 - EEG

#### Participants

29 new volunteers recruited from the UC San Diego community completed Experiment 2. All participants self-reported normal or corrected-to-normal visual acuity and gave both written and oral informed consent as required by the local Institutional Review Board. Each participant was tested in a single 2.5-3 hour experimental session (the exact duration varied across participants depending on the amount of time needed to set up and calibrate the EEG equipment). Unlike Experiment 1, participants were not trained on the categorization task prior to testing. We adopted this approach in the hopes of tracking the gradual emergence of categorical biases during learning. However, many participants learned the task relatively quickly (within 40-60 trials), leaving too few trials to enable a direct analysis of this possibility. Data from one participant were discarded due to a high number of EOG artifacts (over 35% of trials); the data reported here reflect the remaining 28 participants.

#### Behavioral Tasks

In separate runs (where “run” refers to a continuous block of 60 trials lasting approximately 6.5 minutes), participants performed orientation mapping and category discrimination tasks similar to those used in Experiment 1. During both tasks a rapid series of letters (subtending 1.14° × 1.14° from a viewing distance of 55 cm) was presented at fixation, and an aperture of 150 iso-oriented bars (subtending 0.5° × 1.2°) was presented in the periphery. The aperture of bars had inner and outer radii of 1.96° and 9.13°, respectively. On each trial, the bars were assigned one of 15 possible orientations (again 0°-168° in 12° increments) and flickered at a rate of 30 Hz. Each bar was randomly replotted within the aperture at the beginning of each “up” cycle. Letters in the RSVP stream were presented at a rate of 6.67 Hz

During orientation mapping runs, participants detected and reported the presence of a target letter (an X or Y) that appeared at an unpredictable time during the interval from +750 msec to +2250 ms following stimulus onset. Responses were made on a USB-compatible number pad. During category discrimination runs, participants ignored the RSVP stream and instead reported whether the orientation of the bar aperture was an exemplar from category “1” or category “2”. As in Experiment 1, we randomly designated one of the 15 possible stimulus orientations as the category boundary such that the seven orientations counterclockwise to this value were assigned to Category 1 and the seven orientations clockwise to this value were assigned to Category 2. Participants could respond at any point during the trial, but the stimulus was presented for a total of 3000 msec. Trials were separated by a 2.5 – 3.25 sec inter-trial-interval (randomly selected from a uniform distribution on each trial). Each participant completed four (N = 1), five (N = 10), six (N = 8), seven (N = 8), or eight (N = 1) blocks of the category task and three (N = 1), four (N = 1), five (N = 5), six (N = 12), seven (N = 8), or eight (N = 1) blocks of the orientation mapping task.

#### EEG Acquisition and Preprocessing

Participants were seated in a dimly lit, sound-attenuated, and electrically shielded recording chamber (ETS Lindgren) for the duration of the experiment. Continuous EEG was recorded from 128 Ag-AgCl^−^ scalp electrodes via a Biosemi “Active Two” system (Amsterdam, Netherlands). The horizontal electrooculogram (EOG) was recorded from additional electrodes placed near the left and right canthi, and the vertical EOG was recorded from electrodes placed above and below the right eye. Additional electrodes were placed over the left and right mastoids. The horizontal and vertical EOG were recorded from electrodes placed over the left and right canthi and above and below the right eye (respectively). Electrode impedances were kept well below 20 kΩ, and recordings were digitized at 1024 Hz.

After testing, the entire EEG time series at each electrode was high- and low-pass filtered (3^rd^ order zero-phase forward and reverse Butterworth) at 0.1 and 50 Hz and re-referenced to the average of the left and right mastoids. Data from both tasks were epoched into intervals spanning −1000 to +4000 msec from stimulus onset; the relatively large pre- and post-stimulus epochs were included to absorb filtering artifacts that could affect later analyses. Trials contaminated by EOG artifacts (horizontal eye movements > 2° and blinks) were identified and excluded from additional analyses. Across participants an average of 5.58% (±1.67%) and 8.74% (±1.84%) of trials from the orientation mapping and category discrimination tasks were discarded (respectively). Finally, noisy channels (those with multiple deflections ≥ 100 µV over the course of the experiment) were visually identified and eliminated (mean number of removed electrodes across participants ±1 S.E.M. = 2.25 ± 0.64).

Next, we identified a set of electrodes-of-interest (EOIs) with strong responses at the stimulus’ flicker frequency (30 Hz). Data from each task were re-epoched into intervals spanning 0 to 3000 msec around stimulus onset and averaged across trials and tasks (i.e., RSVP and category discrimination), yielding a *k* electrode by *t* sample data matrix. We computed the evoked power at the stimulus’ flicker frequency (30 Hz) by applying a discrete Fourier transform to the average time series at each electrode and selected the 32 electrodes with the highest evoked power at the stimulus’ flicker frequency for further analysis. These electrodes were typically distributed over occipitoparietal electrode sites (see Figure 12).

To isolate stimulus-specific responses, the epoched timeseries at each electrode was resampled to 256 Hz and then bandpass filtered from 29 to 31 Hz (zero-phase forward and reverse 3^rd^ order Butterworth). We next estimated a set of complex Fourier coefficients describing the power and phase of the 30 Hz response by applying a Hilbert transformation to the filtered data. To visualize and quantify orientation-selective signals from frequency-specific responses, we first constructed a complex-valued data set B_1_(t) (*m* electrodes × *n_train_* trials). We then estimated a complex-valued weight matrix W(t) (*m* channels × *k* filters) using B_1_(t) and a basis set of idealized orientation-selective filters C_1_. Finally, we estimated a complex-valued matrix of channel responses C_2_(t) (*m* channels × *n_test_* trials) given W(t) and complex-valued test data set B_2_(t) (*m* electrodes × *n_test_* trials) containing the complex Fourier coefficients measured during the category discrimination task. Trial-by-trial and sample-by-sample response functions were shifted in the same manner described above so that category biases would manifest as a rightward (clockwise) shift towards the center of Category 2. We estimated the evoked (i.e., phase-locked) power of the response at each filter by computing the squared absolute value of the average complex-valued coefficient for each filter after shifting. Categorical biases were quantified using the same curve fitting analysis described in the main text.

To obtain an unbiased estimate of orientation selectivity in each electrode, we ensured that the training data set B_1_(t) contained an equal number of trials for each stimulus orientation (0-168° in 12° increments). For each participant, we identified the stimulus orientation θ with the *N* fewest repetitions in the orientation mapping data set after EOG artifact removal. Next, we constructed the training data set B_1_(t) by randomly selecting (without replacement) 1:*N* trials for each stimulus orientation. Data from this training set were used to estimate a set of orientation weights for each electrode and these weights were in turn used to estimate a response for each hypothetical orientation channel during the category discrimination task. To ensure that our method generalized across multiple combinations of orientation mapping trials, we repeated this analysis 100 times and averaged the results across permutations.

### Experiment 3 - EEG

#### Participants

8 volunteers recruited from the Florida Atlantic University community completed Experiment 3. All participants self-reported normal or corrected-to-normal visual acuity and gave both written and oral informed consent as required by the local Institutional Review Board. Each participant was tested in a single 2-2.5 hour experimental session (the exact duration varied across participants depending on the amount of time needed to set up and calibrate the EEG equipment).

#### Behavioral Tasks

Participants performed six blocks of a spatial recall task followed by multiple blocks of a delayed match-to-category (DMC) task. Both tasks used identical stimulus and display geometry. During the spatial recall task, participants were shown a sample display containing a disc (diameter 2.5° from a viewing distance of 60 cm) rendered in one of 12 polar locations (0° to 330° in 30° increments) along the perimeter of an imaginary circle centered at fixation (radius 7.5°). The sample display was shown for 250 ms and followed by a 1750 ms blank delay. At the end of each trial, participants were shown a mouse cursor and instructed to click on the position of the disc shown in the sample display. Participants were instructed to prioritize accuracy over speed, though a 3000 ms response deadline was imposed. Each trial was followed by a 1500-2200 ms blank interval (randomly sampled from a uniform distribution on each trial). Each block featured 72 trials (six repetitions per stimulus position) and lasted approximately six minutes. EEG data recorded during this task were used to train a position-specific inverted encoding model (see below). Each participant completed six blocks of this task.

After completing the spatial recall task, participants performed a delayed match-to-category task. Participants were shown stimuli in the same 12 positions used during the spatial recall task. However, for each participant we defined a category boundary such that half of the possible stimulus positions were assigned membership in Category 1 and the remaining half were assigned membership in Category 2. For example, the category boundary could be set such that positions [315, 345, 15, 45, 75, 105] comprised Category 1 while positions [135, 165, 195, 225, 255, 285] comprised Category 2. The location of the category boundary was randomly and independently chosen for each participant and held constant throughout the experiment. At the beginning of each trial, a sample disc appeared in one of the 12 possible stimulus locations for 250 ms. After a 1750 ms delay period, a probe disc was presented. The probe could occupy any of the 11 stimulus positions not occupied by the sample, and participants were required to judge whether the position of the probe matched the category of the sample stimulus via keypress. Participants were instructed to prioritize accuracy over speed, but a 3000 ms response limit was imposed. Feedback (correct vs. incorrect) was presented at the end of each trial. Participants completed 5 (N = 1) or 8 (N = 7) blocks of 72 trials.

#### EEG Acquisition and Preprocessing

Continuous EEG was recorded from 63 Ag/Ag-Cl^−^ scalp electrodes via a Brain Products actiCHamp amplifier. An additional electrode was placed over the right mastoid. Data were recorded with a right mastoid reference and later re-referenced to the algebraic mean of the left and right mastoids (10-20 site TP9 served as the left mastoid reference). The horizontal and vertical electrooculogram (EOG) was recorded from electrodes placed on the left and right canthi and above and below the right eye, respectively. All electrode impedances were kept below 15 kΩ, and recordings were digitized at 1000 Hz. Recorded data were bandpass filtered from 1 to 50 Hz (3^rd^ order zero-phase forward and reverse Butterworth filters), epoched from a period spanning −1000 to +3000 ms relative to the start of each trial, and baseline corrected from −250 to 0 ms. Muscle and electrooculogram artifacts were removed from the data using independent components analysis (ICA) as implemented in EEGLAB (Delorme & Makeig, 2004). Reconstructions of stimulus locations were computed from the spatial topography of induced alpha-band (8-12 Hz) power measured across 17 occipitoparietal electrode sites: O1, O2, Oz, PO7, PO3, POz, PO4, PO8, P7, P5, P3, P1, Pz, P2, P4, P6, and P8.

#### Inverted Encoding Model

Experiment 3 relied on a fundamentally different signal than Experiment 2 (induced-alpha-band activity vs. evoked 30 Hz power, respectively). Following earlier research (Kok et al., 2017; Ester et al., 2018; Nouri & Ester, 2019), we used a variant of the IEM approach described in Experiment 2 to compute location channel responses. We first isolated alpha-band activity, by bandpass filtering the raw EEG time series at each electrode from 8-12 Hz (zero-phase forward and reverse filters as implemented by EEGLAB’s “eegfilt” function), yielding a real-valued signal *f*(*t*). The analytic representation of *f*(*t*) was obtained by applying a Hilbert transformation:

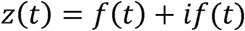

where 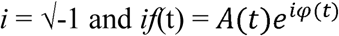. Induced alpha power was computed by extracting and squaring the instantaneous amplitude *A*(*t*) of the analytic signal *z*(*t*). We modeled alpha power at each scalp electrode as a weighted sum of 12 location-selective channels, each with an idealized tuning curve (a half-wave rectified cosine raised to the 12^th^ power). The maximum response of each channel was normalized to 1, thus units of response are arbitrary. The predicted responses of each channel during each trial were arranged in a *k* channel by *n* trials design matrix *C*. Separate design matrices were constructed to track the locations of the blue and red discs across trials (i.e., we reconstructed the locations of the blue and red discs separately, then later sorted these reconstructions according to cue condition). The relationship between the data and the predicted channel responses *C* is given by a general linear model of the form:

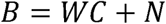

where B is a *m* electrode by *n* trials training data matrix, W is an *m* electrode by *k* channel weight matrix, and *N* is a matrix of residuals (i.e., noise).

To estimate *W*, we constructed a “training” data set containing an equal number of trials from each stimulus location (i.e., 45-360° in 45° steps) condition. We first identified the location φ with the fewest *r* repetitions in the full data set after EOG artifact removal. Next, we constructed a training data set *B_trn_* (*m* electrodes by *n* trials) and weight matrix *C_trn_* (*n* trials by *k* channels) by randomly selecting (without replacement) 1:*r* trials for each of the eight possible stimulus locations (ignoring cue condition; i.e., the training data set contained a mixture of neutral and valid trials). The training data set was used to compute a weight for each channel *C_i_* via least-squares estimation:

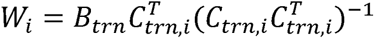

where *C_trn,i_* is an *n* trial row vector containing the predicted responses of spatial channel *i* during each training trial.

After estimating the weight matrix *W*, we next estimated a set of spatial filters *V* that capture the underlying channel responses while accounting for correlated variability between electrode sites (i.e., the noise covariance; Kok et al. 2017):

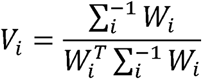

where *Σ_i_* is the regularized noise covariance matrix for channel *i* and estimated as:

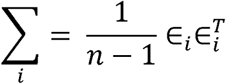

where *n* is the number of training trials and *ε_i_* is a matrix of residuals:

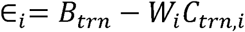

Estimates of ε*_i_* were obtained by regularization-based shrinkage using an analytically determined shrinkage parameter (see Blankertz et al. 2011; Kok et al. 2017). An optimal spatial filter *v_i_* was estimated for each channel *C_i_*, yielding an *m* electrode by *k* filter matrix *V*. Next, we constructed a “test” data set *B_tst_* (*m* electrodes by *n* trials) containing data from all trials not included in the training data set and estimated trial-by-trial channel responses *C_tst_* (*k* channels × *n* trials) from the filter matrix *V* and the test data set:

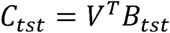

Trial-by-trial channel responses were interpolated to 360°, circularly shifted to a common center (0°, by convention), and sorted by category membership. As in Experiments 1 and 2A, reconstructions were shifted and aligned so that any bias would manifest as a shift toward Category B (clockwise). Finally, to ensure internal reliability this entire analysis was repeated 50 times, and unique (randomly chosen) subsets of trials were used to define the training and test data sets during each permutation. The results were then averaged across permutations.

#### Eye Movement Control Analyses – Experiments 2 and 3

Systematic biases in eye position can contribute to orientation and location performance (e.g., Quax et al., 2019). We did not collect eye position data from Experiment 1 (fMRI). However, different tasks were used to train and test the encoding model, which can be an effective way of mitigating the effects of eye movements on stimulus decoding (Mostert et al., 2018). We also collected electrooculogram (EOG) data during Experiments 2 and 3 (EEG). To examine whether eye position varied as a function of stimulus position during these experiments, we regressed trial-by-trial horizontal EOG recordings (in µV) onto the orientation of a to-be-categorized stimulus (Experiment 2) or the location of a to-be-categorized disc (Experiment 3). In both experiments, we identified and excluded trials contaminated by large horizontal EOG artifacts (≥ 40 µV, which corresponds to a horizontal displacement of 2.5° assuming a voltage threshold of 16 µV per degree; Lins et al., 1993), but smaller variations in eye positions – for example, along the inner stimulus aperture – may have escaped detection. Using Experiment 2 as an example, we considered two possibilities. First, participants may have foveated the inner aperture of the stimulus at a polar location matching its orientation. To illustrate, participants could foveate the inner aperture of a 45° stimulus at a polar angle of 45° or 225°; likewise, they could foveate the inner aperture of a 168° stimulus at a polar angle of 168° or 348°. Second, participants may have foveated the inner aperture of each stimulus matching the center of the category it belonged to. We tested these possibilities by calculating predicted horizontal eye positions under the assumption that participants foveated the inner stimulus aperture at locations matching its orientation or the center of the relevant category. Specifically, we converted records of stimulus orientation (or the center of the category to which the stimulus belonged) to polar format and scaled the resulting estimates by the radius of the inner stimulus aperture, then regressed these estimates onto horizontal EOG activity (in µV). If there is a systematic relationship between eye position and either stimulus orientation or category at any point during a trial, then this analysis should yield regression coefficients reliably greater than 0 µV. Identical analyses were used to examine systematic relationships between horizontal eye position and stimulus location in Experiment 3.

## Results

### Experiment 1 - fMRI

We trained eight human volunteers to categorize a set of orientations into two groups, Category 1 and Category 2. The stimulus space comprised a set of 15 oriented stimuli, spanning 0-168° in 12° increments (Figure 1A-B). For each participant, we randomly designated one of these 15 orientations as a category boundary such that the seven orientations anticlockwise to the boundary were assigned membership in Category 1 and the seven orientations clockwise to the boundary were assigned membership in Category 2. Each participant completed a one-hour training session prior to scanning. Each participant’s category boundary was kept constant across all behavioral training and scanning sessions. Many participants self-reported that they learned the rule delineating the categories in one or two 5-minute blocks of trials. Consequently, task performance measured during scanning was extremely high, with errors and slow responses present only for exemplars immediately adjacent to the category boundary (Figure 1C-D). During each scanning session, participants performed the category discrimination task and an orientation model estimation task where they were required to report the identity of a target letter embedded within a rapid stream presented at fixation while a task-irrelevant grating flickered in the background. Data from this task were used to compute an unbiased estimate of orientation selectivity for each voxel in visual areas V1-hV4v/V3A (see below).

We first examined whether category training increased the similarity of activation patterns evoked by exemplars from the same category (i.e., acquired similarity). We tested this by training a linear decoder (support vector machine) to discriminate between activation patterns associated with exemplars at the center of each category (48° from the boundary), then used the trained classifier to predict the category membership of exemplars immediately adjacent to the category boundary (±12°; Figure 2A). This analysis was performed separately for the orientation mapping and category discrimination tasks. We reasoned that if category training homogenizes activation patterns evoked by members of the same category, then decoding performance should be superior during the category discrimination task relative to the orientation mapping task. This is precisely what we observed (Figure 2B). For example, near-boundary decoding performance in V1 was reliably above chance during the category discrimination task (p < 0.0001, false-discovery-rate-corrected bootstrap test), but not during the orientation mapping task when the category boundary was irrelevant and the oriented stimulus was unattended (p = 0.38). Importantly, the absence of robust decoding performance during the orientation mapping task cannot be attributed to poor signal, as a decoder trained and tested on activation patterns associated with exemplars at the center of each category (Figure 2C) yielded above-chance decoding during both behavioral tasks (Figure 2D; M = 0.58 and 0.69 for the mapping and discrimination tasks, respectively; *p* < 0.01, bootstrap test). Collectively, these results suggest that category training can alter population-level responses at very early stages of the visual processing hierarchy.

**Figure 2.**
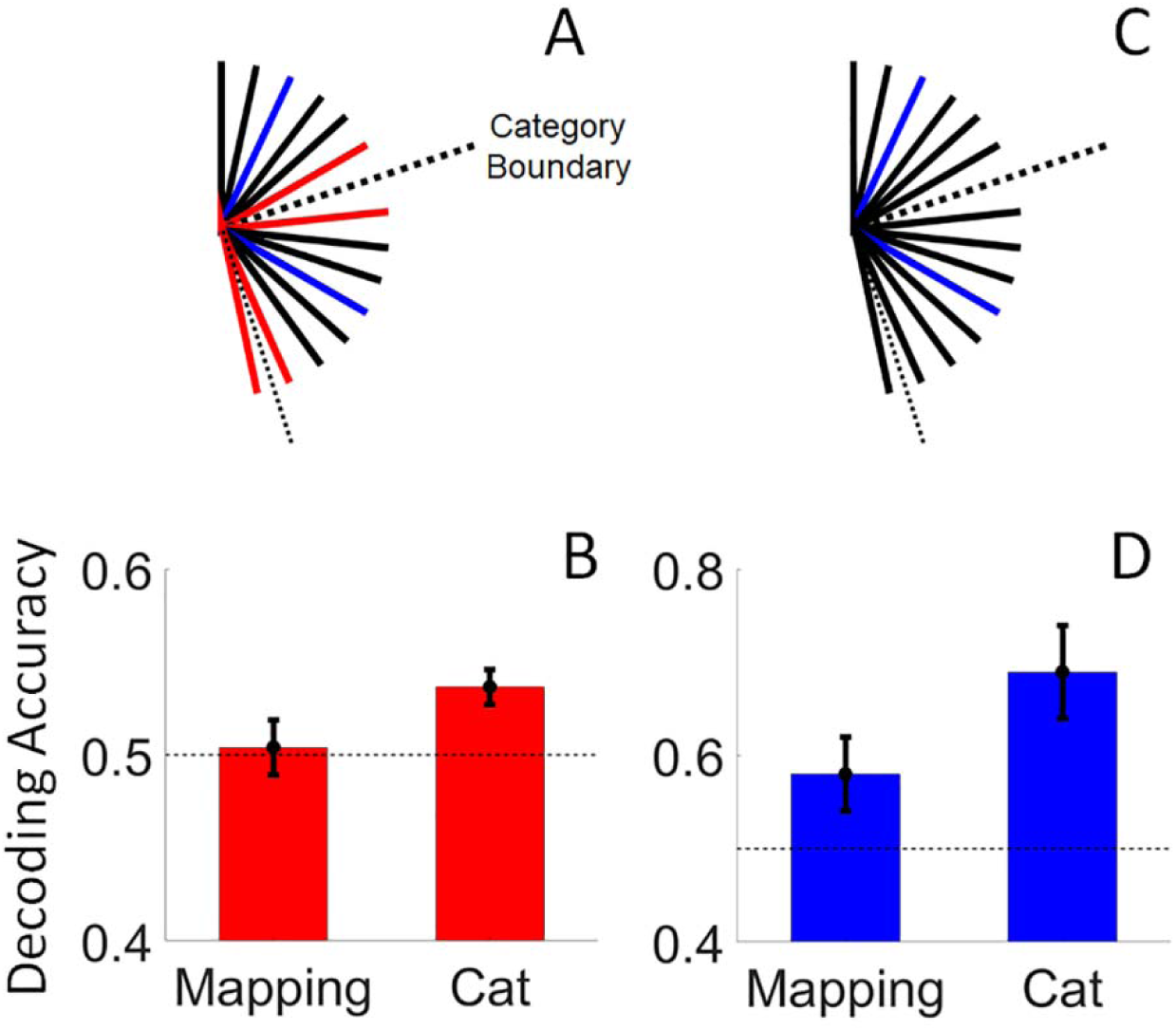
Category Decoding Performance. (A) We trained classifiers on activation patterns evoked by exemplars at the center of each category boundary during the orientation mapping and category discrimination task (blue lines), then used the trained classifier to predict the category membership of exemplars adjacent to the category boundary (red lines). (B) Decoding accuracy was significantly higher during the category discrimination task relative to the orientation mapping task (p = 0.01), suggesting that activation patterns evoked by exemplars adjacent to the category boundary became more similar to activation patterns evoked by exemplars at the center of each category during the categorization task. The absence of robust decoding performance during the orientation mapping task cannot be attributed to poor signal or a uniform enhancement of orientation representations by attention, as a decoder trained and tested on activation patterns associated with exemplars at the center of each category (C) yielded above-chance decoding during both behavioral tasks (D). Decoding performance was computed from activation patterns in V1. Error bars depict ±1 S.E.M.

To better understand how category training influences orientation-selective activation patterns in early visual cortical areas, we used an inverted encoding model (Brouwer & Heeger, 2011) to generate model-based reconstructed representations of stimulus orientation from these patterns. For each visual area (e.g., V1), we first modelled voxel-wise responses measured during the orientation mapping task as a weighted sum of idealized orientation channels, yielding a set of weights that characterize the orientation selectivity of each voxel (Figure 3A). In the second phase of the analysis, we reconstructed trial-by-trial representations of stimulus orientation by combining these weights with the observed pattern of activation across voxels measured during each trial of the category discrimination task, resulting in single-trial reconstructed channel response function that contains a representation of stimulus orientation for each ROI on each trial (Figure 3B). Finally, we sorted trial-by-trial reconstructions according to category membership such that any bias would manifest as a clockwise (rightward) shift of the orientation representation towards the center of Category 2 and quantified biases towards this category using a curve-fitting analysis (Methods).

**Figure 3.**
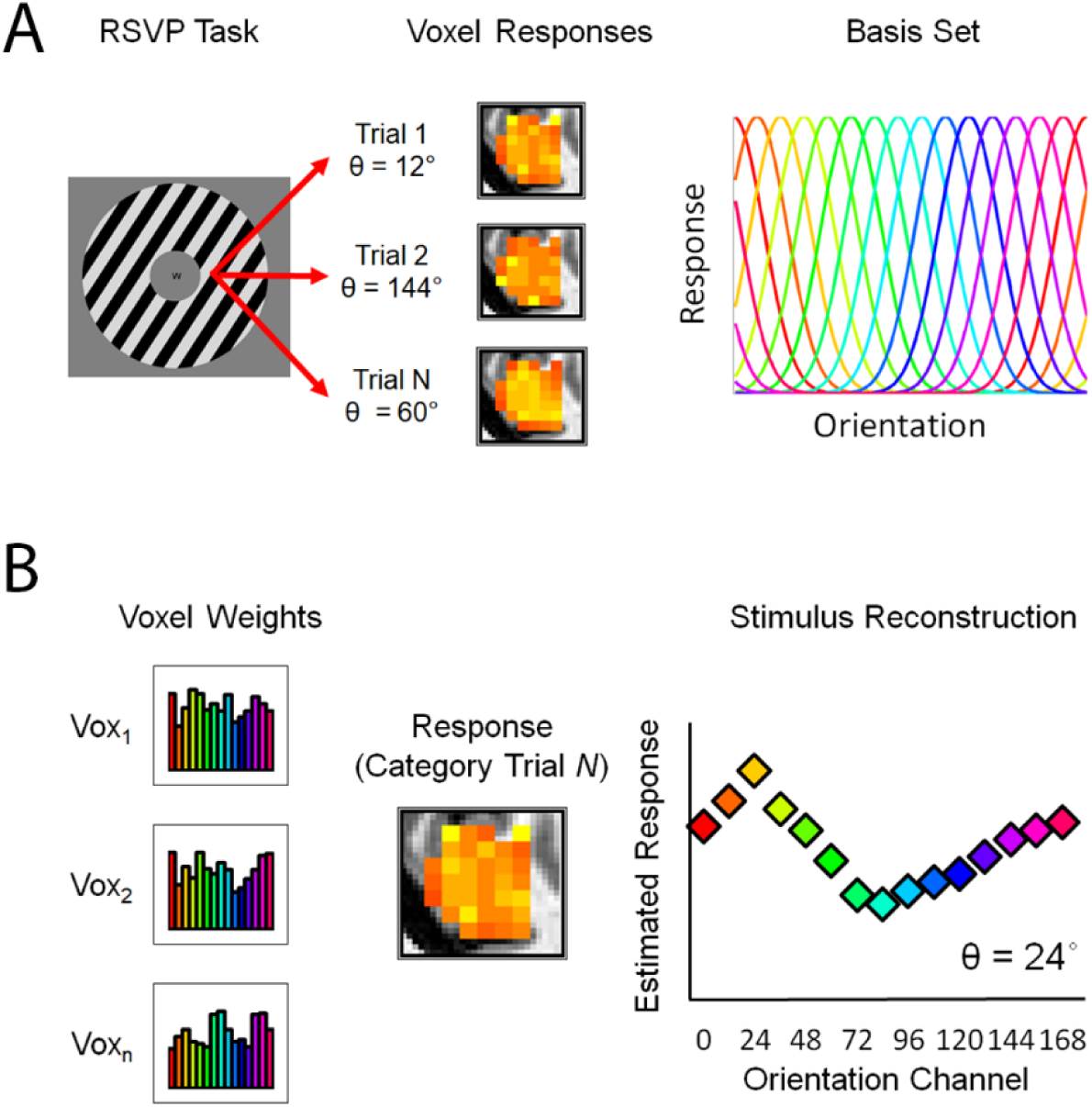
Inverted Encoding Model. (A) In the first phase of the analysis, we estimated an orientation selectivity profile for each voxel retinotopically organized V1-hV4/V3a using data from an independent orientation mapping task. Specifically, we modeled the response of each voxel as a set of 15 hypothetical orientation channels, each with an idealized response function. (B) In the second phase of the analysis, we computed the response of each orientation channel from the estimated orientation weights and the pattern of responses across voxels measured during each trial of the category discrimination task. The resulting reconstructed channel response function (CRF) contains a representation of the stimulus orientation (example; 24 deg), which we quantified via a curve-fitting procedure.

Note that stimulus orientation was irrelevant during the orientation mapping task used for model weight estimation. We therefore reasoned that voxel-by-voxel responses evoked by each oriented stimulus would be largely uncontaminated by its category membership. Indeed, the logic of our analytical approach rests on the assumption that orientation-selective responses are quantitatively different during the orientation mapping and category discrimination tasks: if identical category biases are present in both tasks then the orientation weights computed using data from either task will capture that bias and reconstructed representations of orientation will not exhibit any category shift. This is precisely what we observed when we used a cross-validation approach to reconstruct stimulus representations separately for the orientation mapping and category discrimination tasks (Figure 4).

**Figure 4.**
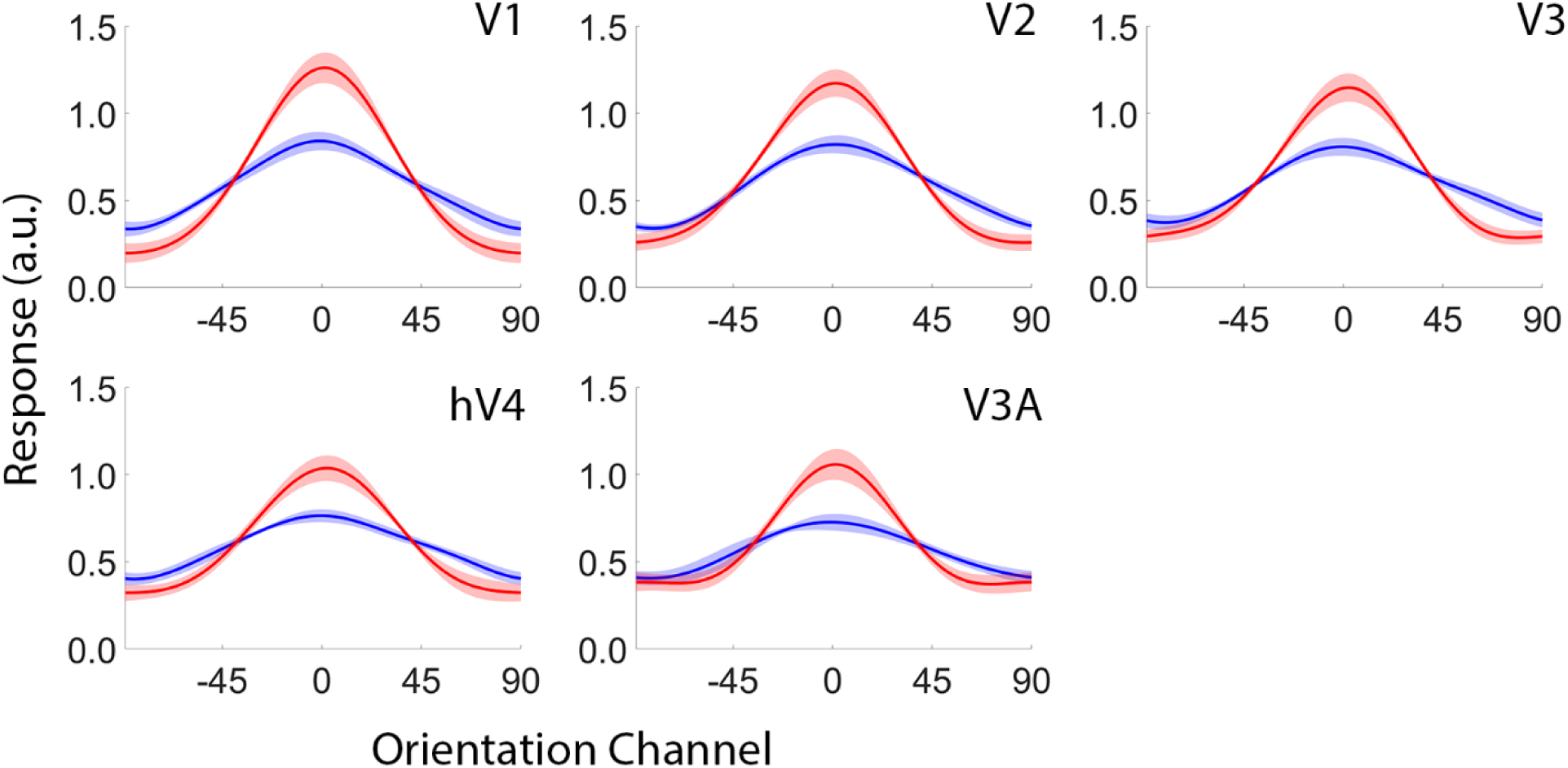
Reconstructions of stimulus orientation during the orientation mapping task (blue) and the category discrimination task (red). Reconstructions were computed using a leave-one-run-out cross validation approach where data from N-1 runs were used to estimate channel weights and data from the remaining run were used to estimate channel responses. This procedure was iterated until all runs had been used to estimate channel responses and the results were averaged over permutations. No categorical biases were observed in any visual area for either task. Shaded regions depict ±1 within-participant S.E.M. a.u., arbitrary units.

As shown in Figure 5, reconstructed representations of orientation in visual areas V1, V2, and V3 were systematically biased away from physical stimulus orientation and towards the center of the appropriate category (shifts of 22.13°, 26.65°, and 34.57°, respectively; *P* < 0.05, bootstrap test, false-discovery-rate [FDR] corrected for multiple comparisons across regions; see Figure 6 for separate reconstructions of Category 1 and Category 2 orientations and Figure 7 for participant-by-participant reconstructions plotted by visual area). Similar, though less robust biases were also evident in hV4v and V3A (mean shifts of 9.73° and 6.45°, respectively; *p* > 0.19). A logistic regression analysis established that categorical biases in V1-V3 were strongly correlated with variability in overt category judgments (Figure 8). That is, trial-by-trial category judgments were more strongly associated with the responses of orientation channels near the center of each category rather than those near the physical orientation of the stimulus. Importantly, because the location of the boundary separating categories 1 and 2 was randomly selected for each participant, it is unlikely that categorical biases shown in Figure 5 reflect intrinsic biases in stimulus selectivity in early visual areas (e.g., due to oblique effects; Sun et al., 2013).

**Figure 5.**
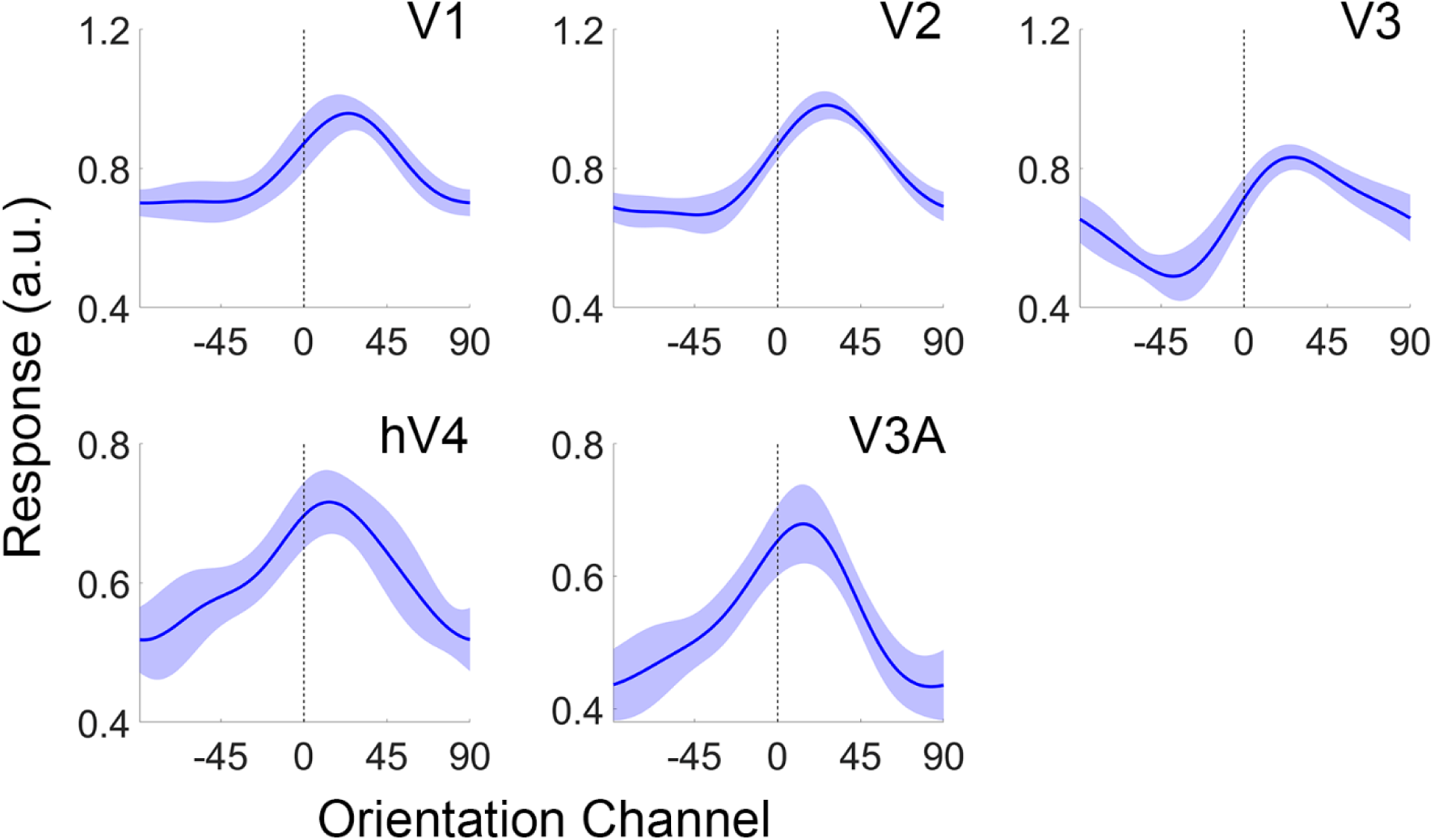
Reconstructed representations of Orientation in Early Visual Cortex. The vertical bar at 0° indicates the actual stimulus orientation presented on each trial. Channel response functions (CRFs) from Category 1 and Category 2 trials have been arranged and averaged such that any categorical bias would manifest as a clockwise (rightward) shift in the orientation representation towards the center of Category B. Shaded regions are ±1 within-participant S.E.M (see Methods). Note change in scale between visual areas V1-V3 and hV4-V3A. a.u., arbitrary units.

**Figure 6.**
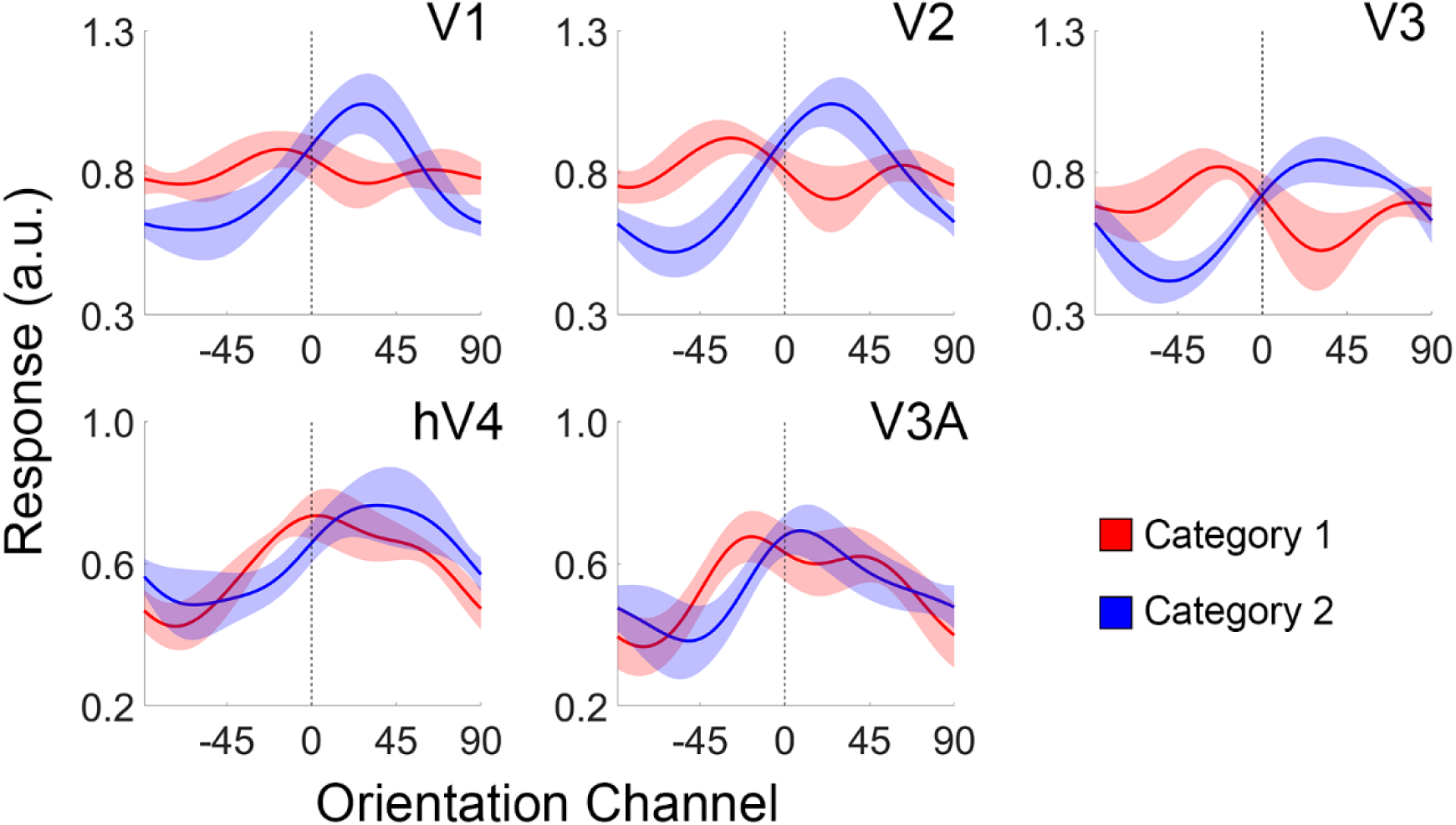
Stimulus Reconstructions during Category 1 and Category 2 trials. Shaded regions are ±1 within-participant S.E.M. a.u., arbitrary units.

**Figure 7.**
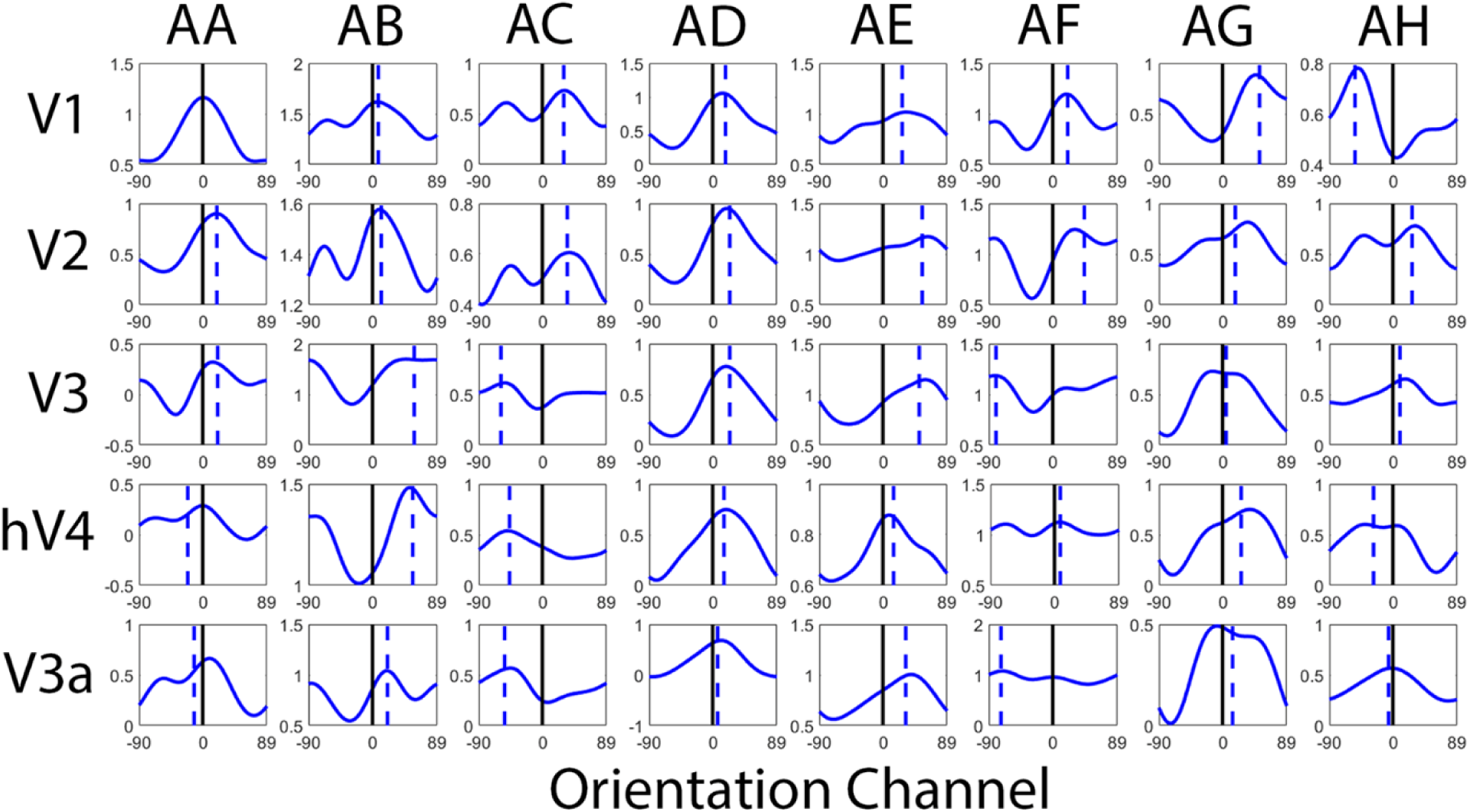
Participant-level Stimulus Reconstructions. Each panel plots a reconstructed representation of stimulus orientation for a given participant (columns) and visual area (rows). Dashed blue lines are the estimated peak of each reconstruction (obtained via curve-fitting). Ordinate units are arbitrary.

**Figure 8.**
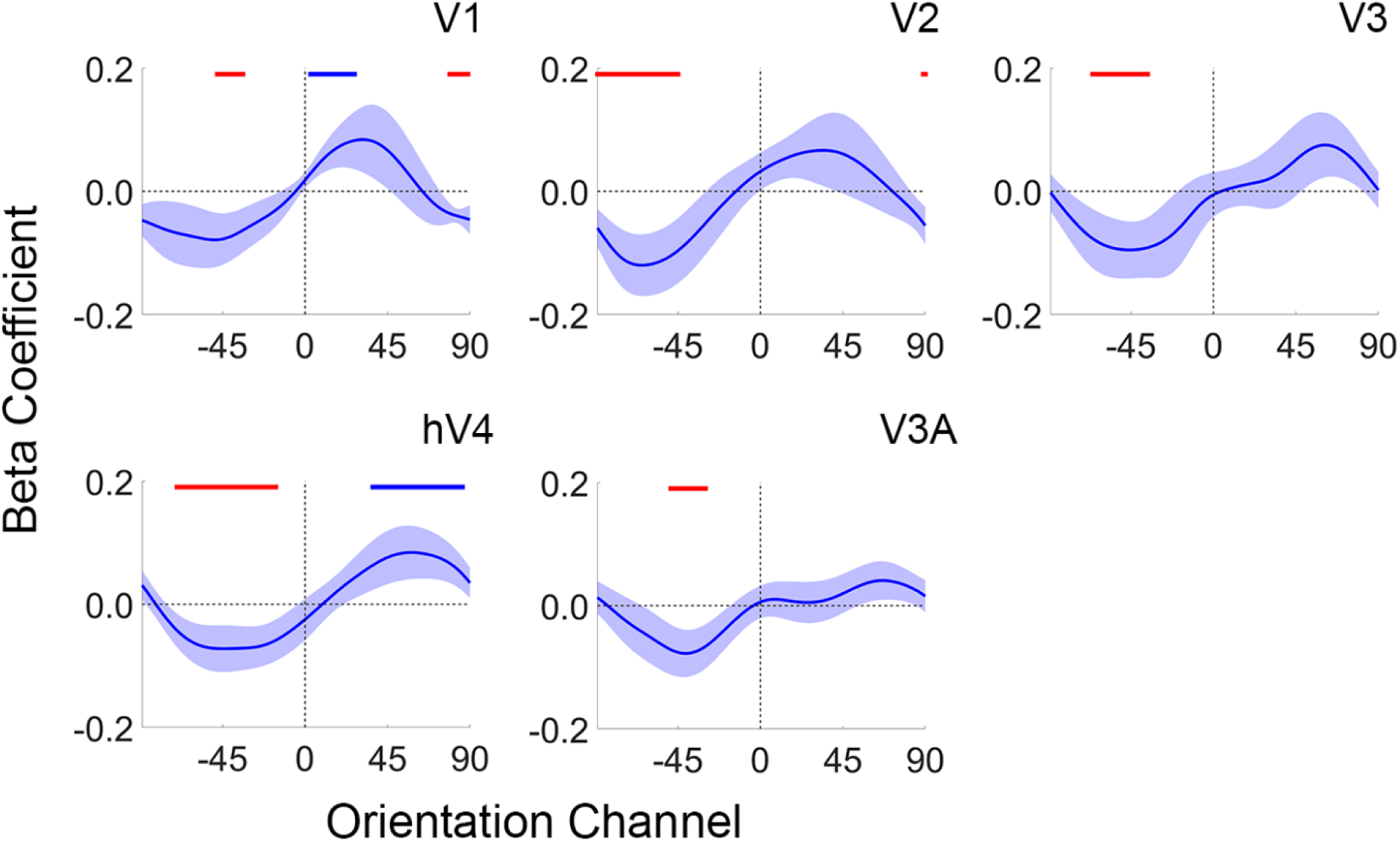
Categorical Biases predict Choice Behavior. Each plot shows a logistic regression of each orientation channel’s response onto trial-by-trial variability in category judgments. A positive coefficient indicates a positive relationship between an orientation channel’s response and the correct category judgment (i.e., Category B), while a negative coefficient indicates a negative relationship between an orientation channel’s response and correct category judgment (i.e., Category A). Red and blue horizontal lines at the top of each plot depict orientation channels whose estimated β coefficients are significantly below or above zero, respectively (FDR-corrected permutation test; p < 0.05). Shaded regions are ±1 within-participant S.E.M.

The category biases shown in Figure 5 may be the result of an adaptive process that facilitates task performance by enhancing the discriminability of physically similar but categorically distinct stimuli. Consider a hypothetical example where an observer is tasked with discriminating between two physically similar exemplars on opposite sides of a category boundary. Discriminating between these alternatives should be challenging as each exemplar evokes a similar and highly overlapping response pattern. However, discrimination performance could be improved if the responses associated with each exemplar are made more separable via acquired distinctiveness following training (or equivalently, an acquired similarity between exemplars adjacent to the category boundary and exemplars near the center of each category). Similar changes would be less helpful when an observer is tasked with discriminating between physically and categorically distinct exemplars, as each exemplar already evokes a dissimilar and non-overlapping response. From these examples, a simple prediction can be derived: categorical biases in reconstructed representations of orientation should be largest when participants are shown exemplars adjacent to the category boundary and progressively weaker when participants are shown exemplars further away from the category boundary.

We tested this possibility by sorting stimulus reconstructions according to the angular distance between stimulus orientation and the category boundary (Figure 9). As predicted, reconstructed representations of orientations adjacent to the category boundary were strongly biased by category membership, with larger biases for exemplars nearest to the category boundary (µ = 42.62°, 24.16°, and 20.12° for exemplars located 12°, 24°, and 36° from the category boundary, respectively; FDR-corrected bootstrap p < 0.0015), while reconstructed representations of orientations at the center of each category exhibited no signs of bias (µ = - 3.98°, p = 0.79; the direct comparison of biases for exemplars adjacent to the category boundary and in the center of each category was also significant; p < 0.01). Moreover, the relationship between average category bias and distance from the category boundary was well-approximated by a linear trend (slope = −14.38°/step; r^2^ = 0.96). Thus, category biases in reconstructed representation are largest under conditions where they would facilitate behavioral performance and absent under conditions where they would not.

**Figure 9.**
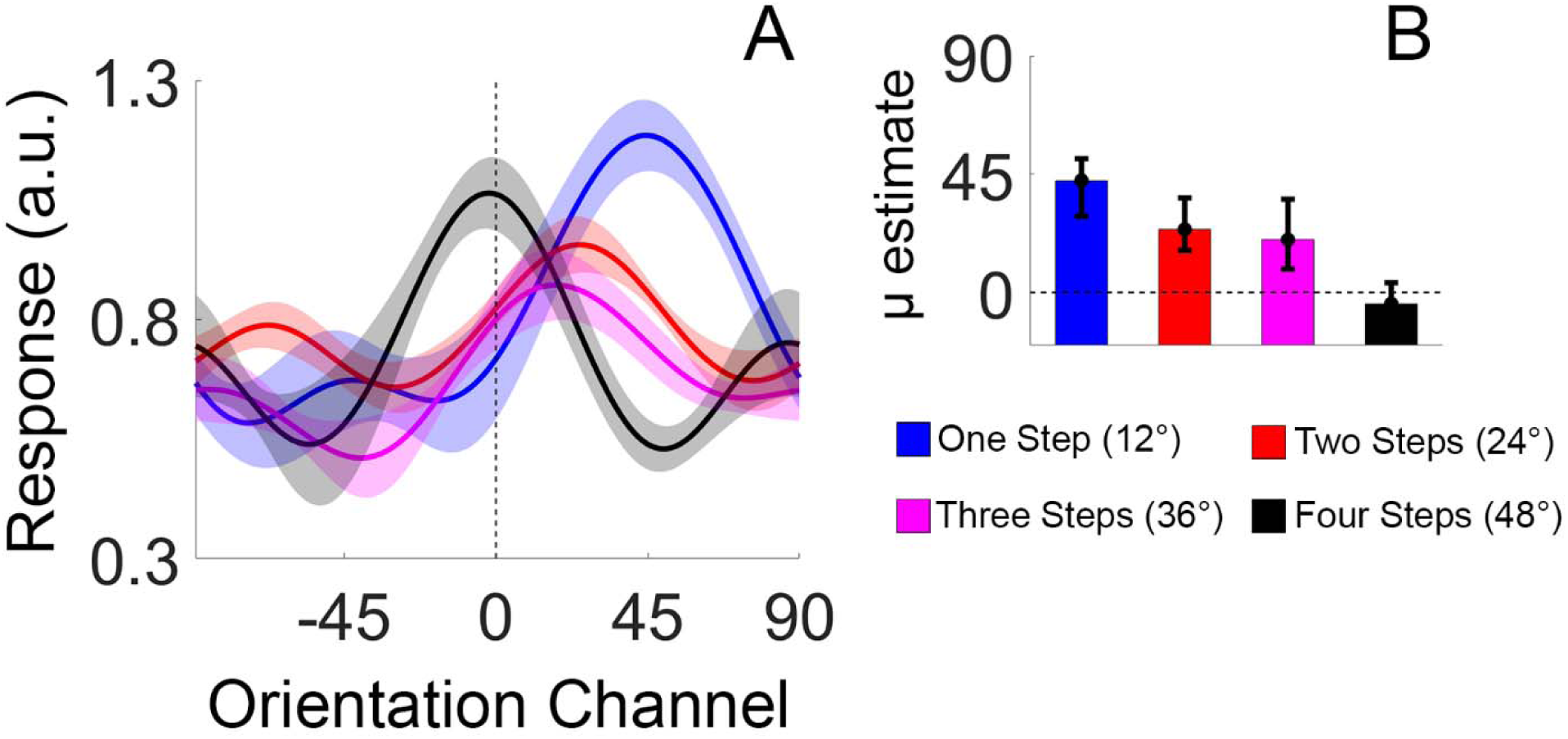
Category Biases Scale Inversely with Distance from the Category Boundary. (A) The reconstructions shown in Fig. 3 sorted by the absolute angular distance between each exemplar and the category boundary. In our case, the 15 orientations were bisected into two groups of 7, with the remaining orientation serving as the category boundary. Thus, the maximum absolute angular distance between each orientation category and the category boundary was 48°. Participant-level reconstructions were pooled and averaged across visual areas V1, V2, and V3 as no differences were observed across these regions. Shaded regions are ±1 within-participant S.E.M. (B) shows the amount of bias for exemplars located 1, 2, 3, or 4 steps from the category boundary (quantified via a curve-fitting analysis). Error bars are 95% confidence intervals. a.u., arbitrary units.

Category-selective signals have been identified in multiple brain areas, including portions of lateral occipital cortex, inferotemporal cortex, posterior parietal cortex, and lateral prefrontal cortex (Sigala & Logothetis, 2002; Freedman et al., 2011; Freedman & Assad, 2006; Folstein et al., 2012; Davis & Poldrack, 2013; Pourtois et al., 2008; Mack et al., 2013). We identified category selective information in many of these same regions using a whole-brain searchlight-based decoding analysis where a classifier was trained to discriminate between exemplars from Category 1 and Category 2 (independently of stimulus orientation; Figure 10 and Methods). Next, we used the same inverted encoding model described above to reconstruct representations of stimulus orientation from activation patterns measured in each area during each of the orientation mapping and category discrimination tasks (reconstructions were computed using a “leave-one-participant-out” cross-validation routine to ensure that reconstructions were independent of the decoding analysis used to define category-selective ROIs). We were able to reconstruct representations of stimulus orientation in many of these regions during the category discrimination task, but not during the orientation mapping task (where stimulus orientation was task-irrelevant; Figure 11). This is perhaps unsurprising as representations in many mid-to-high order cortical areas are strongly task-dependent (e.g., Silver et al., 2005). As our analytical approach requires an independent and unbiased estimate of each voxel’s orientation selectivity (e.g., during the orientation mapping task), this meant that we were unable to probe categorical biases in reconstructed representations in these regions.

**Figure 10.**
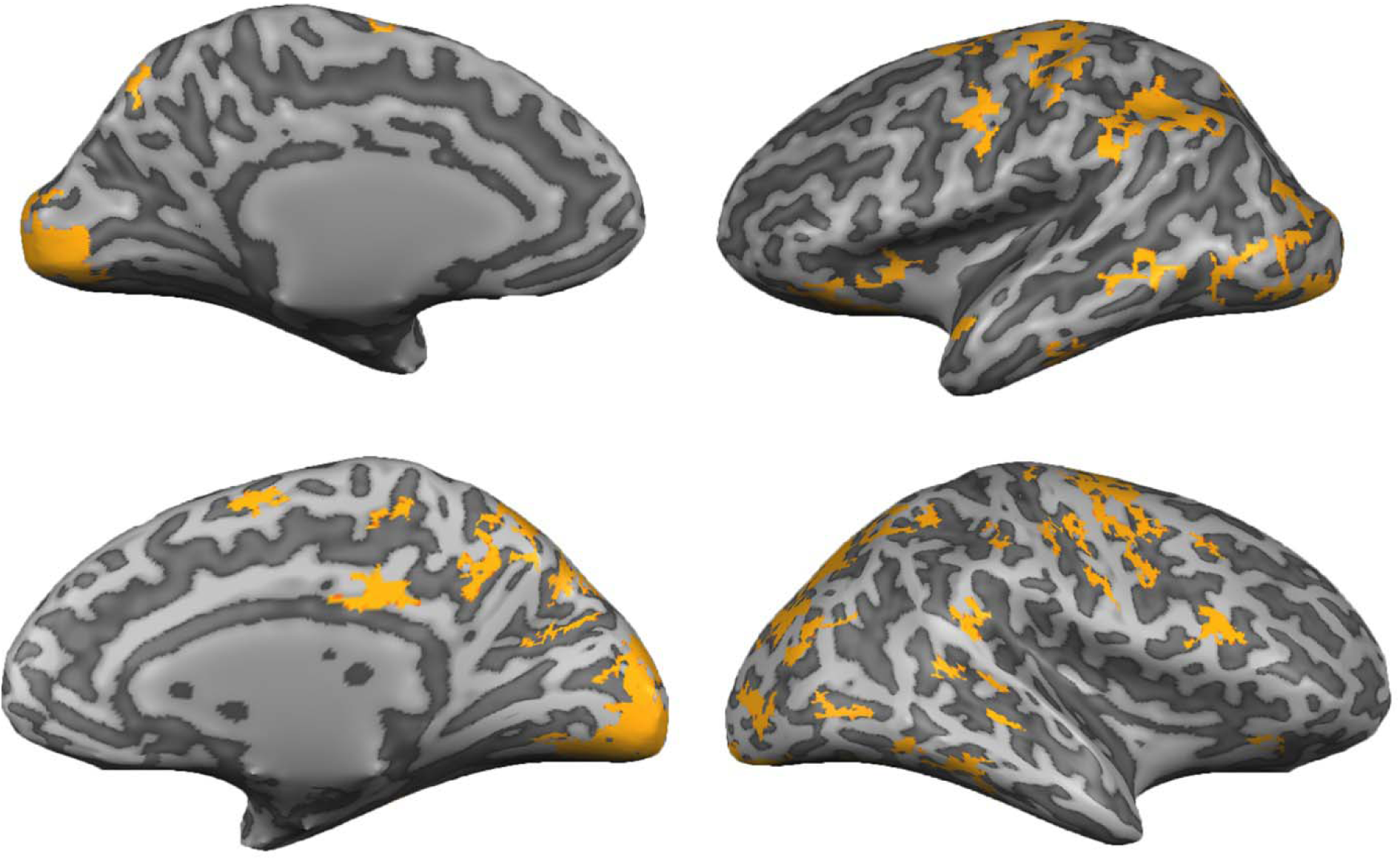
Cortical Areas Supporting Robust Decoding of Category Information. We trained a linear support vector machine to discriminate between activation patterns associated with Category A and Category B exemplars (see *Searchlight Classification Analysis*; Methods). The trained classifier revealed robust category information in multiple visual, parietal, temporal, and prefrontal cortical areas, including many regions previously associated with categorization (e.g., posterior parietal cortex and lateral prefrontal cortex).

**Figure 11.**
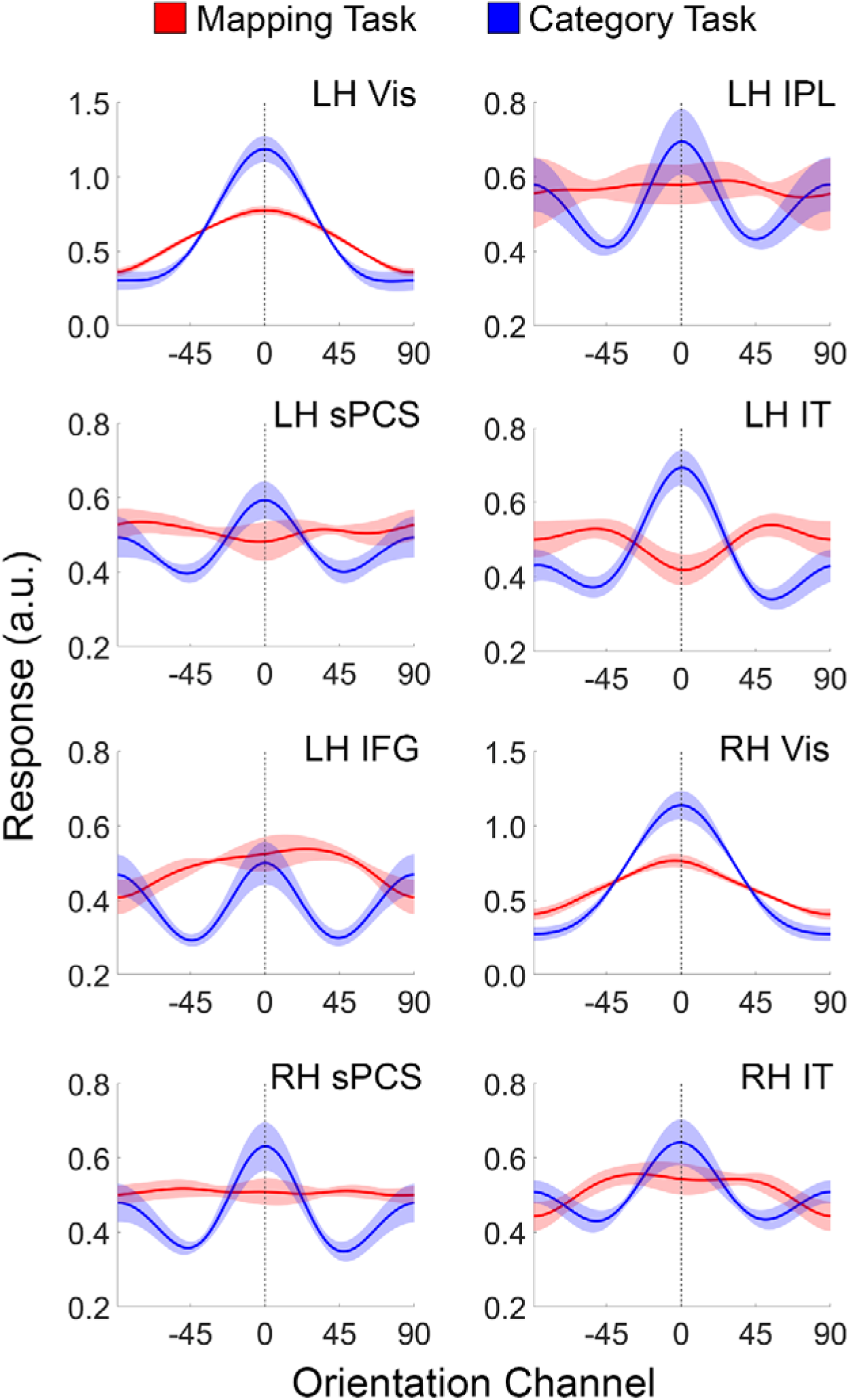
Stimulus Reconstructions in Visual, Parietal, and Frontal cortical areas during the Orientation Mapping and Categorization Tasks. During the orientation mapping task, participants detected and reported the identity of a target presented in a stream of letters at fixation. During the categorization experiment, participants categorized stimulus orientation into two discrete groups. Shaded regions are ±1 within-participant S.E.M. IPL, inferior parietal lobule; IPS, intraparietal sulcus; sPCS, superior precentral sulcus; IT, inferotemporal cortex, IFG, inferior frontal gyrus. a.u., arbitrary units.

### Experiment 2 - EEG

Due to the sluggish nature of the hemodynamic response, the category biases shown in Figures 5 and 9 could reflect processes related to decision making or response selection rather than stimulus processing. In a second experiment, we evaluated the temporal dynamics of category biases using EEG. Specifically, we reasoned that if the biases shown in Figures 5 and 9 reflect processes related to decision making, response selection, or motor planning, then these biases should manifest only during a period shortly before the participants’ response. Conversely, if the biases are due to changes in how sensory neural populations encode features, they should be evident during the early portion of each trial. To evaluate these alternatives, we recorded EEG while a new group of 28 volunteers performed variants of the orientation mapping and categorization tasks used in the fMRI experiment. On each trial, participants were shown a large annulus of iso-oriented bars that flickered at 30 Hz (i.e., 16.67 ms on, 16.67 ms off; Figure 12A). During the orientation mapping task, participants detected and reported the identity of a target letter (an X or a Y) that appeared in a rapid series of letters over the fixation point. Identical displays were used during the category discrimination task, with the caveat that participants were asked to report the category of the oriented stimulus while ignoring the letter stream.

**Figure 12.**
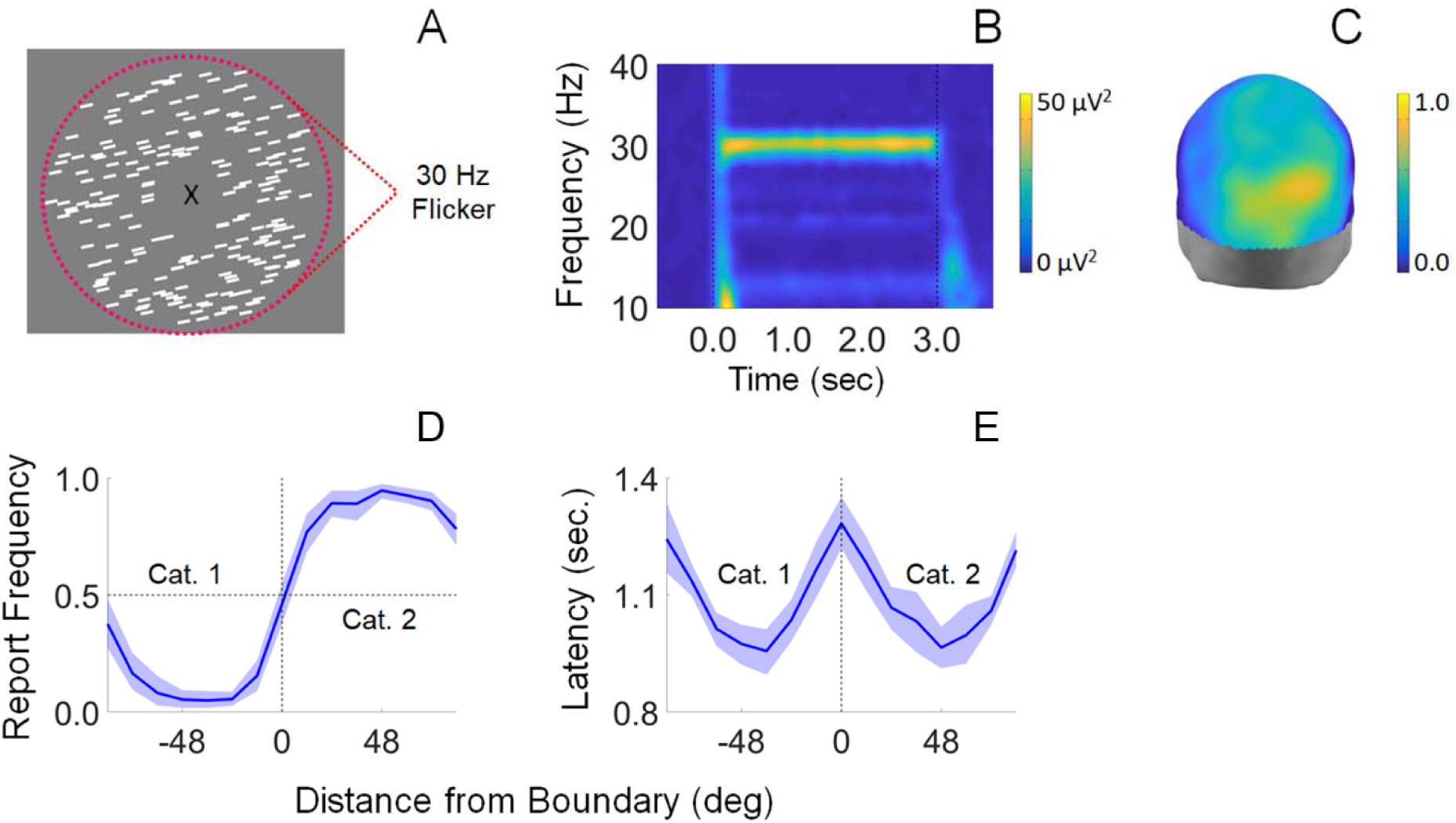
Summary of Experiment 2A. (A) Participants viewed displays containing an aperture of iso-oriented bars flickering at 30 Hz. (B) The 30 Hz flicker entrained a frequency-specific response known as a steady-state visually-evoked potential (SSVEP). (C) Evoked 30 Hz power was largest over occipitoparietal electrode sites. We computed stimulus reconstructions (Fig. 7) using the 32 scalp electrodes with the highest power. The scale bar indicates the proportion of participants (out of 27) for which each electrode site was ranked in the top 32 of all 128 scalp electrodes. (D-E) Participants categorized stimuli with a high degree of accuracy; incorrect and slow responses were observed only for exemplars adjacent to a category boundary. Shaded regions are ±1 within-participant S.E.M.

The 30 Hz flicker of the oriented stimulus elicits a standing wave of frequency-specific sensory activity known as a steady-state visually-evoked potential (SSVEP, Vialatte et al., 2010; Figure 12B). The coarse spatial resolution of EEG precludes precise statements about the cortical source(s) of these signals (e.g., V1, V2, etc.). However, to focus on visual areas (rather than parietal or frontal areas) we deliberately entrained stimulus-locked activity at a relatively high frequency (30 Hz). Our approach was based on the logic that coupled oscillators can only be entrained at high frequencies within small local networks, while larger or more distributed networks can only be entrained at lower frequencies due to conduction delays (Breakspear et al., 2010). Indeed, a topographic analysis showed that evoked 30 Hz activity was strongest over a localized region of occipitoparietal electrode sites. (Figure 12C). As in Experiment 1, participants learned to categorize stimuli with a high degree of accuracy, with errors and slow responses present only for exemplars adjacent to a category boundary (Figure 12D-E)

We computed the power and phase of the 30 Hz SSVEP response across each 3,000 ms trial and then used these values to reconstruct a time-resolved representation of stimulus orientation (Garcia et al., 2013). Our analysis procedure followed that used in Experiment 1: In the first phase of the analysis, we rank-ordered scalp electrodes by 30 Hz power (based on a discrete Fourier transform spanning the 3000 ms trial epoch, averaged across all trials of both the orientation mapping and category discrimination tasks). Responses measured during the orientation mapping task were used to estimate a set of orientation weights for the 32 electrodes with the strongest SSVEP signals (i.e., those with the highest 30 Hz power; see Figure 12C) at each timepoint. In the second phase of the analysis, we used these timepoint-specific weights and corresponding responses measured during each trial of the category discrimination task across all electrodes to compute a time-resolved representation of stimulus orientation (Figure 13A-B). We reasoned that if the categorical biases shown in Figures 5 and 9 reflect processes related to decision making or response selection, then they should emerge gradually over the course of each trial. Conversely, if the categorical biases reflect changes in sensory processing, then they should manifest shortly after stimulus onset. To test this possibility, we computed a temporally averaged stimulus reconstruction over an interval spanning 0 to 250 ms after stimulus onset (Figure 14B). A robust category bias was observed (M = 23.35°; p = 0.014; bootstrap test) suggesting that the intent to categorize a stimulus modulates how neural populations in early visual areas respond to incoming sensory signals.

**Figure 13.**
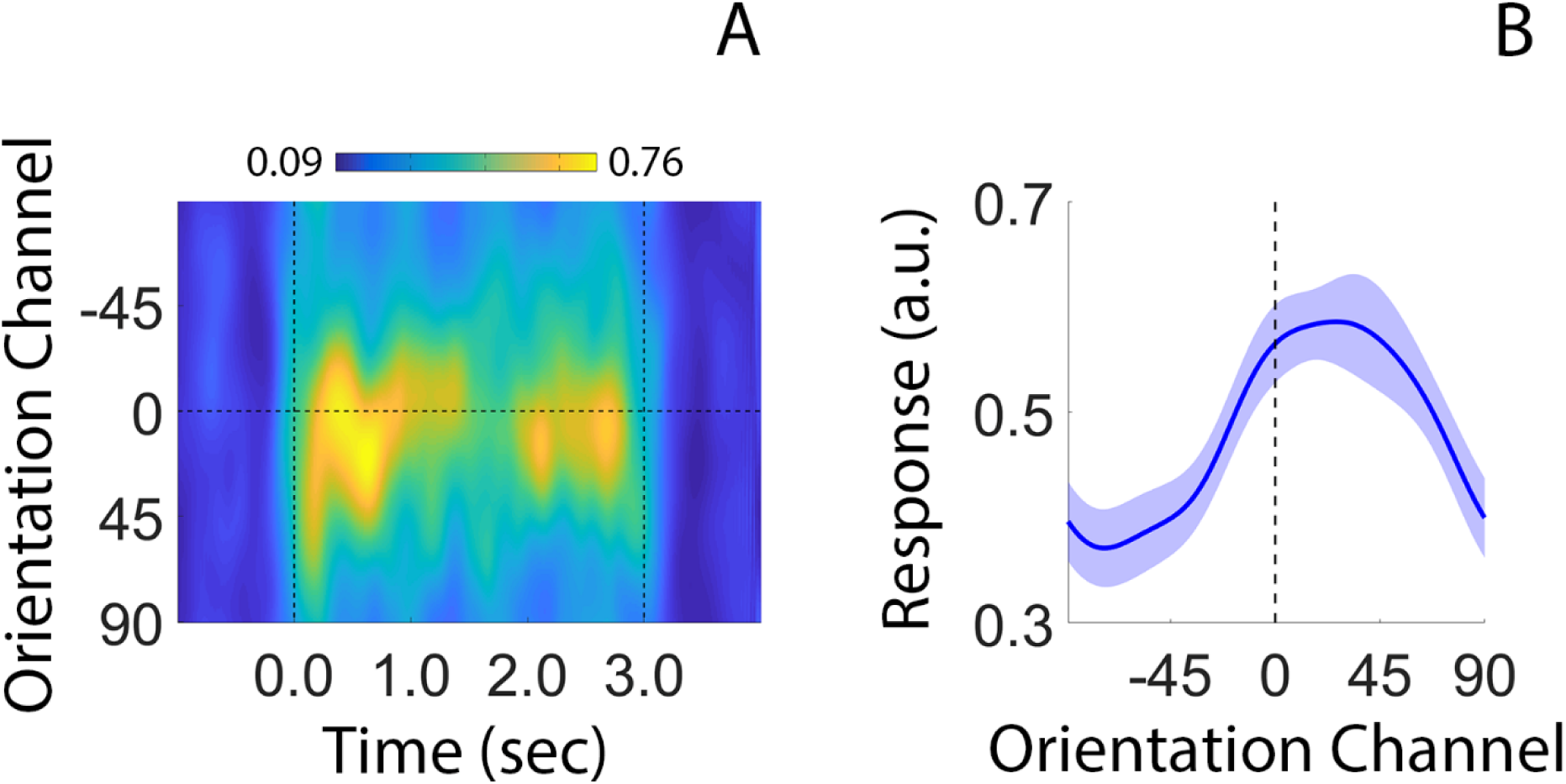
Category Biases Emerge Shortly after Stimulus Onset. (A) Time-resolved reconstruction of stimulus orientation. Dashed vertical lines at time 0.0 and 3.0 seconds mark stimulus on- and offset, respectively. (B) Average channel response function during the first 250 ms of each trial. The reconstructed representation exhibits a robust category bias (*p* < 0.01; bootstrap test). a.u., arbitrary units.

**Figure 14.**
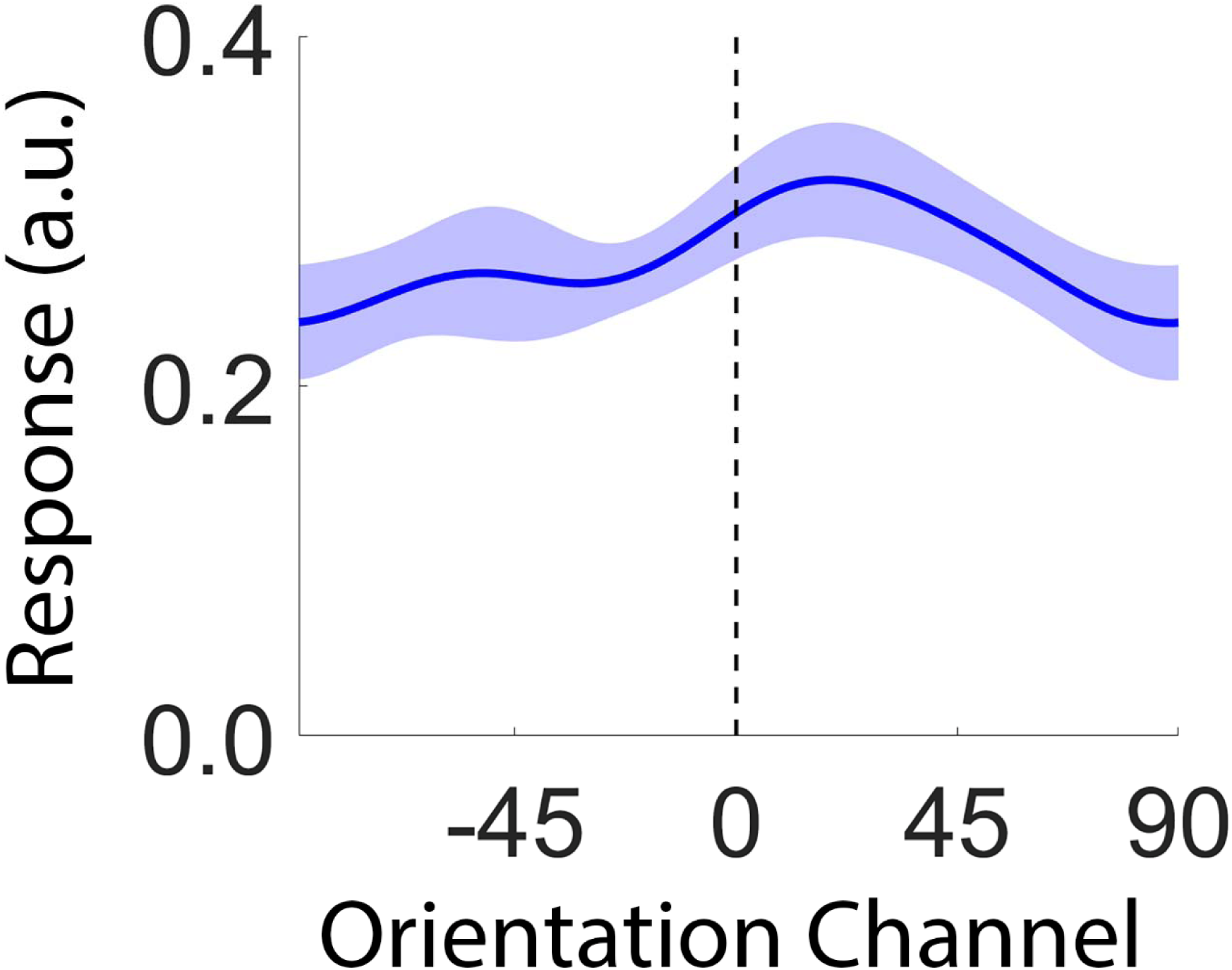
Stimulus- and category information are absent in pre-trial EEG activity. Time-averaged reconstruction computed over an interval spanning −250 to 0 ms relative to stimulus onset. The center of the reconstruction was statistically indistinguishable from 0° (*p* = 0.234; bootstrap test)

Importantly, the bandpass filter used to isolate 30 Hz activity will distort temporal characteristics of the raw EEG signal by some amount. We estimated the extent of this distortion by generating a 3 second, 30 Hz sinusoid with unit amplitude (plus 1 second of pre-and post-signal zero padding) and running it through the same filters used in our analysis path. We then computed the time at which the filtered signal reach 25% of maximum. For an instantaneous filter, this should occur at exactly 1 second (due to the pre- and post-signal zero-padding). We estimated a signal onset of ~877 ms, or 123 ms prior to the start of the signal. Therefore, if reconstruction amplitude is greater than zero at time t, then we can conclude that the pattern of scalp activity used to generate the stimulus reconstruction contained reliable orientation information at time *t* ± 125 ms. The same logic applies to estimates of reconstruction bias as the reconstructions are based on data filtered using the same parameters. Importantly, we also verified that there was no categorical bias in stimulus reconstructions prior to stimulus onset (Figure 14), nor were categorical biases present when we reconstructed stimulus representations using data from the orientation mapping and category discrimination tasks separately (Figure 15).

**Figure 15.**
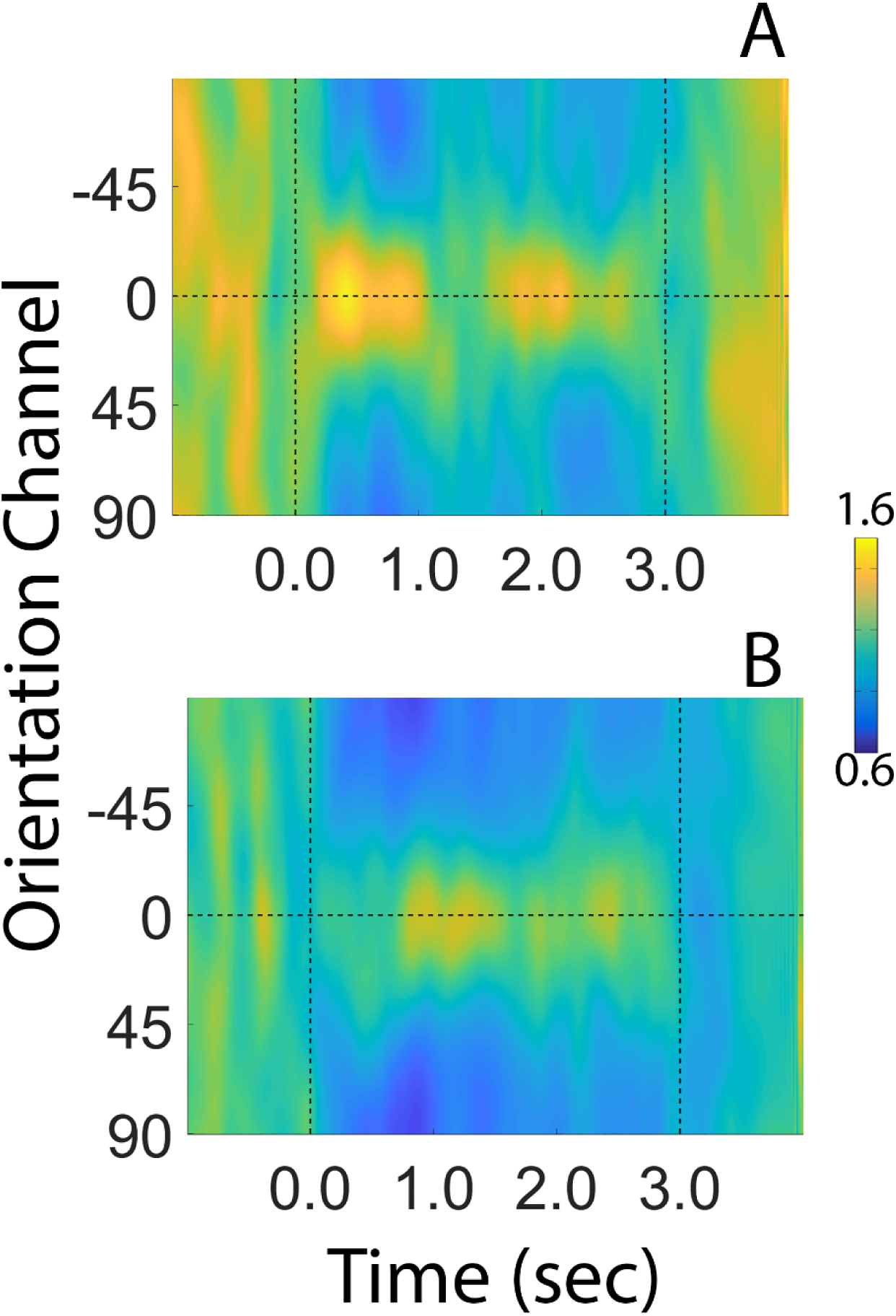
Reconstructions of stimulus orientation during the orientation mapping task (A) and the category discrimination task (B) during Experiment 2. Vertical dashed lines at time 0.0 and 3.0 mark the start and end of each trial, respectively. Reconstructions were computed using a leave-one-run-out cross validation approach where data from N-1 runs were used to estimate channel weights and data from the remaining run were used to estimate channel responses. This procedure was iterated until all runs had been used to estimate channel responses and the results were averaged over permutations. Units of response are arbitrary.

#### Ruling out contributions from eye movements

We identified and removed trials contaminated by large EOG artifacts (blinks and eye movements greater than ~2°). However, small and consistent eye movement patterns could nevertheless contribute to the orientation reconstructions reported here. We examined this possibility by testing whether participants foveated the inner aperture of the stimulus at polar locations matching its orientation (Figure 16A) or at polar locations matching the center of the appropriate category (A vs B; Figure 16B; see Methods for details). No systematic differences in eye position were observed as a function of stimulus orientation or category membership (Figure 16), suggesting that eye movements were not a major contributor to orientation-specific reconstructions.

**Figure 16.**
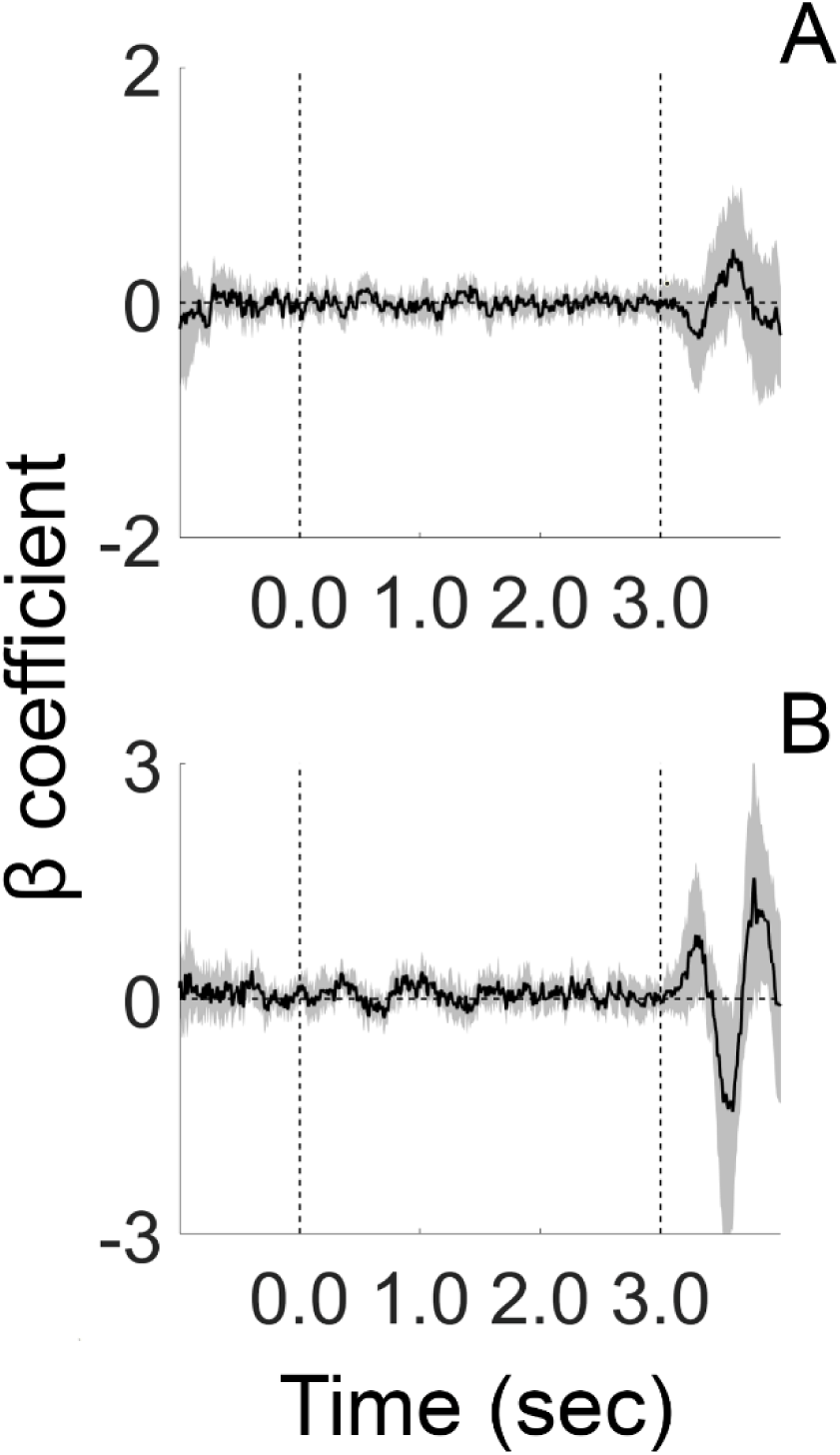
No systematic biases in eye position during orientation categorization (Experiment 2). We regressed trial-by-trial records of stimulus orientation (A) or category (B) onto horizontal EOG activity. Thus, positive coefficients reflect a systematic relationship between stimulus orientation (or category) and eye position. No such biases were observed. Black vertical dashed lines at 0.0 and 3.0 depict the start and end of each trial. Shaded regions depict the 95% within-participant confidence interval of the mean.

### Experiment 3 - EEG

The results of Experiments 1 and 2 suggest that category learning can bias stimulus-specific representations encoded by occipitoparietal cortical areas. However, an alternative explanation is that the biases shown in Figures 5, 9, and 13 reflect mechanisms of response selection or decision making independent of categorical processing. Experiment 3 examined this possibility by examining categorical biases in stimulus-specific memory representations while participants performed a delayed match-to-category (DMC) task. A schematic of the task is shown in Figure 17A-B. At the beginning of each trial a sample disc rendered in one of 12 possible stimulus locations (15-345° polar angle in 30° along the perimeter of an imaginary circle). Half of the disc positions were assigned membership in Category 1, while the remaining half of disc positions were assigned membership in Category 2 (Figure 17A). Participants remembered the position of the sample disc over a blank delay, then judged whether a probe disc was rendered in a position matching the category of the sample disc. The location of the category boundary was randomly determined for each participant, and response feedback (correct vs. incorrect) was provided after every trial. Like Experiment 2, participants were not trained on the DMC task prior to testing and learned to associate specific positions with specific categories through feedback. Before completing the DMC task, participants also completed a spatial working memory task. Display and stimulus geometry were identical during the spatial memory task and the DMC task. On each trial a sample disc was rendered in one of the same 12 positions used during the DMC task. After a short delay, participants recalled the location of the sample disc via mouse click.

**Figure 17.**
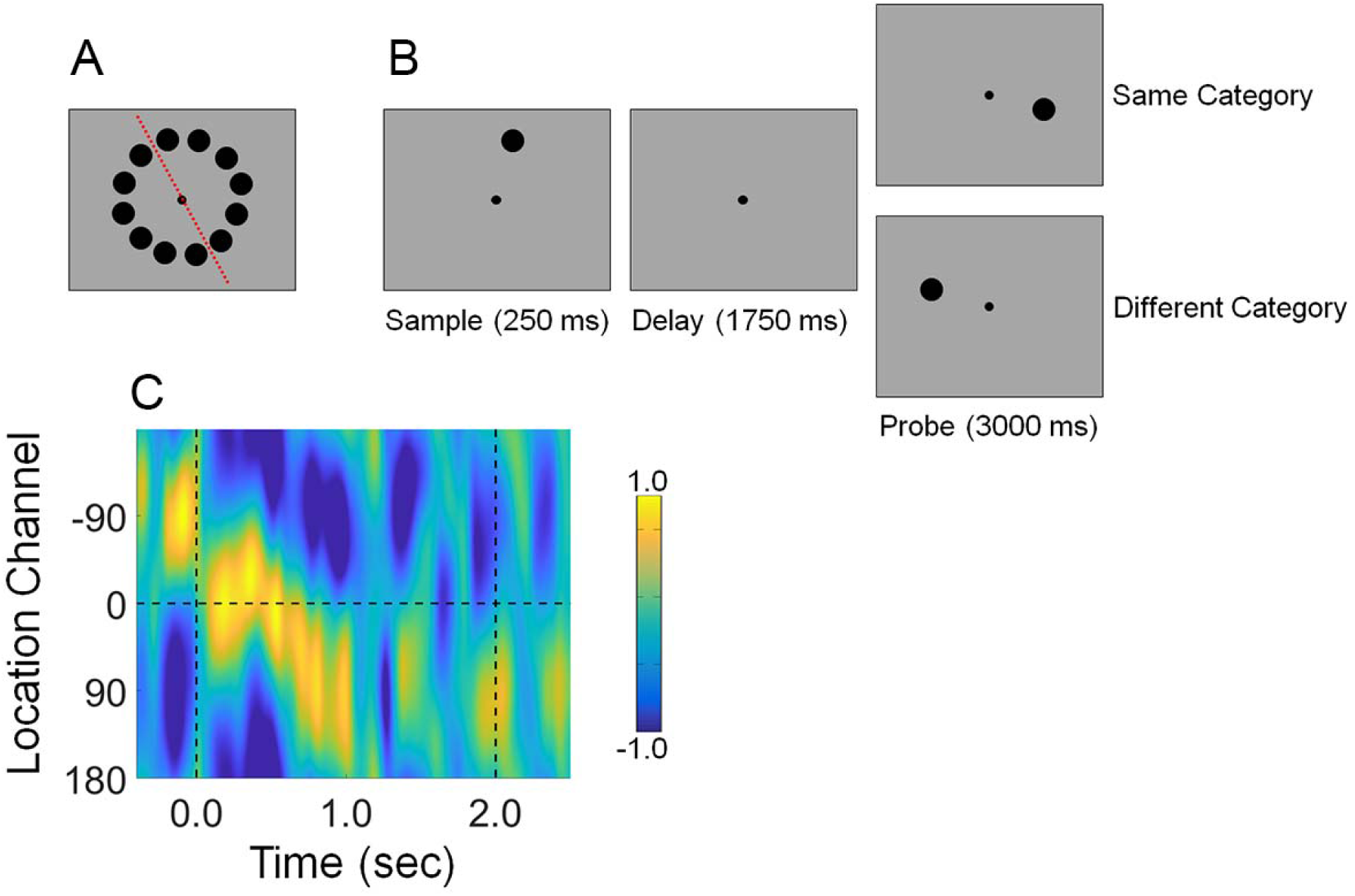
Design and Results of Experiment 3. (A) Possible stimulus locations. The orientation of the category boundary (red dashed line) was randomly determined for each participant (example shown). (B) Delayed match-to-category (DMC) task. Participants remembered the position of a sample disc over a blank delay, then judged whether the location of a probe disc was drawn from the same location category or a different location category. In this example, the categories are defined by the boundary shown in panel A. (C) Location-specific reconstructions computed during the DMC task. Vertical dashed lines at 0.0 and 2.0 sec mark the onset of the sample and probe epochs, respectively. Participants could not prepare a response until the onset of the probe display, yet a robust category bias was observed during the delay period. This suggests that category biases observed in Experiments 1 and 2A are not solely due to mechanisms of response selection.

Following earlier work (e.g., Foster et al., 2016; Samaha et al., 2016; Ester et al., 2018; Nouri & Ester, 2019), we used spatiotemporal patterns of induced alpha-band (8-12 Hz) activity over occipitoparietal electrode sites to track the contents of spatial working memory during the recall and DMC tasks. Specifically, we modeled sample-by-sample estimates of alpha band activity recorded during the spatial recall task as a combination of 12 location filters, each with an idealized tuning curve (a cosine raised to the 12^th^ power). The result of this step is a set of weights that characterizes the location preferences of each scalp electrode. Next, we used these weights and spatiotemporal patterns of alpha-band activity recorded during the DMC task to compute an expected response for each location filter, yielding a time-resolved estimate of stimulus position. Trial-by-trial response functions were shifted to a common center (0° by convention), averaged, and arranged such that any category bias would manifest as a clockwise or positive shift towards the center of Category 2.

As expected, a robust category bias was observed during the delay period of the DMC task (Figure 17C), though unlike Experiment 2 the bias seemed to emerge gradually over the course of the delay period. To quantify this bias, we averaged channel responses from period 0.25 to 2.0 sec after onset of the sample display and fit the resulting function with an exponentiated cosine (*Quantification of Bias in Orientation Representations*, Methods). Mean reconstruction centers were reliably greater than 0° (M = 10.55°; *p* = 0.002, bootstrap test), indicating a robust bias towards the center of the relevant category. Importantly, this bias cannot be explained by mechanisms associated with decision making and response selection: participants could not plan or implement a response until the probe stimulus was presented at the end of the delay period. This result further suggests that the results of Experiments 1 and 2 cannot be wholly explained by mechanisms of response selection or bias.

#### Assessing contributions from eye movements

We identified and removed electrooculogram artifacts from the data via independent components analysis. However, small and consistent eye movement patterns opaque to ICA could nevertheless contribute to the location reconstructions reported here. We examined this possibility by regressing time-resolved estimates of horizontal EOG activity onto remembered stimulus locations. As shown in Figure 18, the regression coefficients linking eye position with remembered locations were indistinguishable from 0 for the duration of each trial, suggesting that eye movements were not a major determinant of location reconstructions.

**Figure 18.**
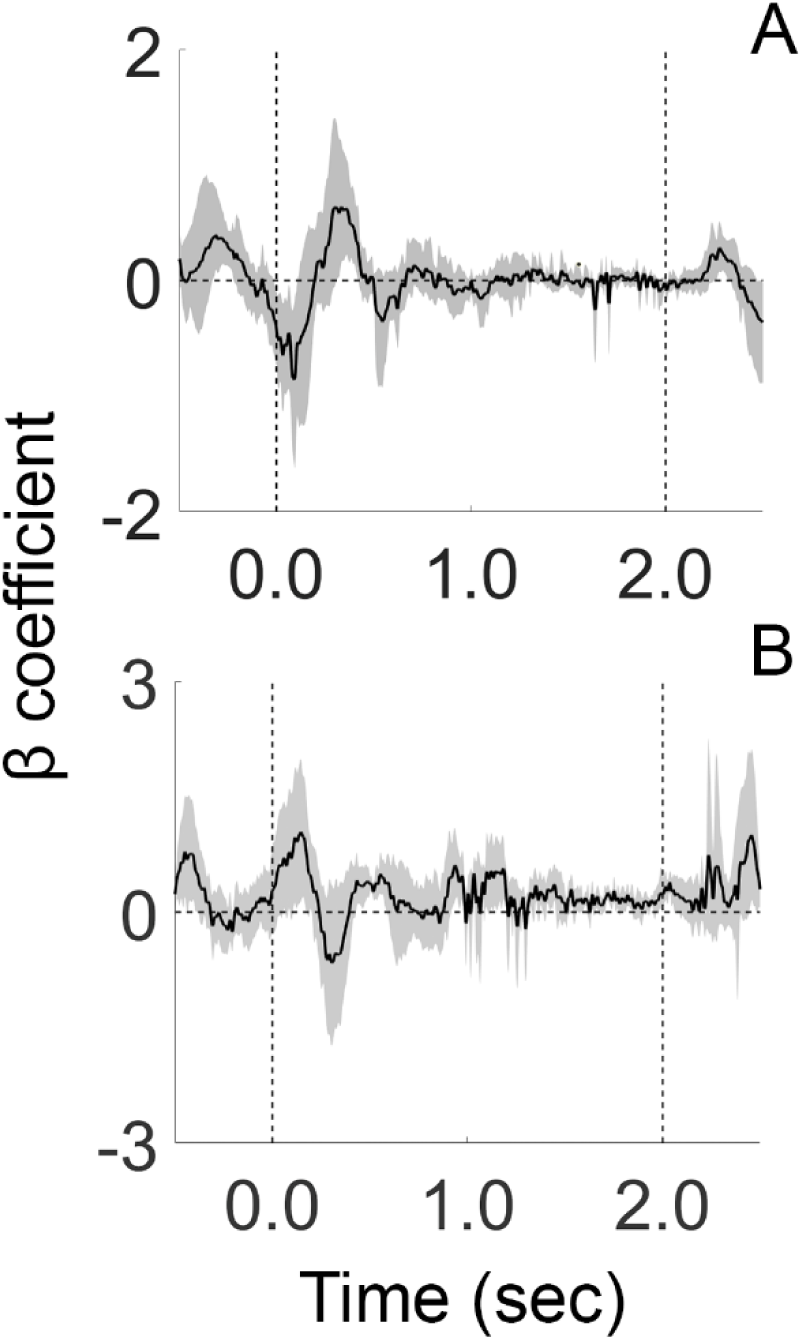
No systematic biases in eye position during location categorization (Experiment 3). We regressed trial-by-trial records of stimulus location (A) or category (B) onto horizontal EOG activity. Thus, positive coefficients reflect a systematic relationship between stimulus orientation (or category) and eye position. No such biases were observed. Black vertical dashed lines at 0.0 and 3.0 depict the start and end of each trial. Shaded regions depict the 95% within-participant confidence interval of the mean.

## Discussion

Our findings suggest that category learning shapes information processing at the earliest stages of the visual system. The results of Experiment 1 showed that representations of a to-be-categorized stimulus encoded by population-level activity in early visual cortical areas were systematically biased by their category membership. These biases were correlated with overt category judgments and were largest for exemplars adjacent to the category boundary. The results of Experiments 2 and 3 demonstrate that similar biases are present in orientation- and location-specific reconstructions computed by human scalp EEG data, and further suggest that our findings cannot be explained by response bias, motor planning, or eye movements.

The categorical biases reported here are strongly task dependent, and therefore must reflect changes in responses caused by transient top-down factors rather than long-term changes in feature or location selectivity. However, the effects of these top down modulations are fundamentally different from task-dependent modulations reported elsewhere. In one example, Ester et al. (2016) asked participants to attend the orientation or luminance of a peripheral grating and found both multiplicative and additive enhancements of orientation-specific reconstructions during the attend orientation condition relative to the attend luminance condition, but no evidence for a shift like the one reported here. In a different study, Byers and Serences (2014) examined changes in orientation-specific reconstructions before and after participants underwent extensive training (10 1-hour sessions) in a challenging orientation discrimination task. We observed changes in the amplitude (i.e., signal-to-noise ratio) of orientation-specific reconstructions following training, but no evidence for a shift like the one reported in the current study. In other studies, Scolari et al. (2012; 2014) examined changes in orientation-specific reconstructions when participants performed fine-grained and coarse-grained orientation discrimination tasks. Participants viewed two oriented gratings in sequence and judged whether they were identical. During one experiment participants were cued to how the second grating might differ from the first (clockwise vs. counterclockwise rotation), while in a second experiment they were not. During the fine-grained discrimination task, the authors observed a modest shift in orientation-specific reconstructions towards “off-target” neural populations that maximally discriminated between two oriented stimuli, but only when participants were cued to expect a clockwise or counterclockwise rotation. While this type of modulation is desirable while performing a fine-discrimination task, it is qualitatively different than the shifts we report in the current experiment, as participants have no way of anticipating what orientation will be presented on each trial, nor the difference between that orientation and the category boundary. Moreover, the shifts reported by Scolari et al. (2012) during fine discriminations were relatively modest – at most few degrees. We report an opposite pattern of findings, where shifts are largest for oriented exemplars immediately adjacent to the category boundary. Thus, while other studies have documented task-dependent changes in orientation-specific reconstructions, those studies have failed to reveal shifts in reconstructed representations (Ester et al., 2016; Byers & Serences 2014) or have revealed modest shifts that follow different patterns from those reported here (Scolari et al. 2012).

Several mechanisms may be responsible for our findings. One possibility is that the orientation preferences of single-units (or populations of units) are systematically shifted towards the center of each category during the category discrimination task, much in the same way that neurons in the rodent auditory system exhibit emergent selectivity for categorically different stimuli (e.g., Xin et al., 2019) or in the same way that the spectral preferences of neural populations are biased by feature-based attention (David et al., 2008; Cukur et al., 2012). These shifts are relatively small at the single unit level but could be amplified by a read-out mechanisms that integrate the responses of large neural populations. A second possibility is that participants strategically apply gain to neural populations that maximally discriminate between to-be-categorized exemplars during the category discrimination task. Here there are no changes in the spectral preferences of single units, but the responses of neurons that respond to stimuli near the category boundary are amplified. These alternatives are not mutually exclusive; nor is this an exhaustive list. Our data cannot resolve these possibilities. For example, several different patterns of single-unit gain changes and/or tuning shifts can produce identical responses in a single fMRI voxel, and different patterns of single-voxel modulation could produce categorical biases in multivariate stimulus reconstructions (see, e.g., Sprague et al., 2018 for a detailed discussion of this issue). Ultimately, targeted experiments that combine non-invasive measurements of brain activity with careful psychophysical measurements and detailed model simulations will be needed to conclusively identify the mechanisms responsible for the category biases we have reported here.

Our findings appear to conflict with results from nonhuman primate research which suggests that sensory cortical areas do not encode categorical information. However, there is reason to suspect that mechanisms of category learning might be qualitatively different in human and non-human primates. For example, our participants learned to categorize stimuli based on performance feedback after approximately 10 minutes of training. In contrast, macaque monkeys typically require six months or more of training using a similar feedback scheme to reach a similar level of performance, and this extensive amount of training may influence how neural circuits code information (e.g., Itthipurripat et al., 2017; Birman & Gardner, 2015). Moreover, there is growing recognition that the contribution(s) of sensory cortical areas to performance on a visual task are highly susceptible to recent history and training effects (Itthipurripiat et al., 2017, Chen et al., 2016; Liu & Pack, 2017). In one example (Liu & Pack, 2017), extensive training was associated with a functional substitution of human visual area V3a for MT+ in discriminating noisy motion patches. Thus, training effects may help explain why previous electrophysiological experiments have found category-selective responses in association but not sensory cortical areas.

Studies of categorization in non-human primates have typically employed variants of a delayed match to category task, where monkeys are shown a sequence of two exemplars separated by a blank delay interval and asked to report whether the category of the second exemplar matches the category of the first exemplar. The advantage of this task is that it allows experimenters to decouple category-selective signals from activity related to decision making, response preparation, and response execution. However, this same advantage also precludes examinations of whether and/or how top-down category-selective signals interact with bottom-up stimulus-specific signals. We made no effort to decouple category-selective and decision-related signals in Experiments 1-2, and thus the category biases observed in those studies could reflect mechanisms of decision making, response selection, or motor planning. The results of Experiment 3 conflict with this interpretation by demonstrating that robust category biases are present during the memory period of a delayed match-to-category task (Freedman & Assad, 2006).

Previous studies have identified cortical modules selective for faces (Kanwisher et al., 1997), locations (Epstein & Kanwisher, 1998), actions (Astafiev et al., 2004; Huth et al., 2012), bodies (Downing et al., 2001); animacy (Konkle & Caramazza, 2013) and size (Konkle & Caramazza, 2013). Other category distinctions (e.g., tools vs. cars) lack specialized processing modules but can be decoded from multivoxel patterns in multiple occipitotemporal regions (e.g., Folstein et al., 2012). Our findings complement these studies by demonstrating that learning a novel and arbitrary category rule is correlated with rapid and reversible changes in stimulus-specific information processing at even earlier stages of the cortical visual processing hierarchy, including V1 (see also Brouwer & Heeger, 2009; 2013). Category-dependent changes in early visual areas may contribute to more complex forms of category selectivity exhibited by upstream cortical areas, including portions of lateral occipital and inferotemporal cortex. This possibility awaits further scrutiny.

To summarize, we have shown that learning a novel and arbitrary category rule based on a simple visual feature (orientation or location) correlates with rapid and reversible changes in sensory and mnemonic representations encoded by regions in early occipitoparietal cortex. These changes correlate with participants’ overt category judgments, are largest for exemplars adjacent to a category boundary, and cannot be explained by decision making or motor preparation. Collectively, these results provide novel and important evidence suggesting that category learning induces rapid-yet-reversable changes in information processing at early stages of the cortical visual processing hierarchy.

## Acknowledgments

Funding provided by NIH R01 EY025872 and a James S. McDonnell Foundation award to J.T.S.. E.F.E., conceived and designed the experiment, collected and analyzed the data, and wrote the paper. T.C.S. provided conceptual input during all phases of the project and edited the paper. J.T.S. supervised all phases of the project. The authors thank Kelvin Lam for assistance with data collection for Experiment 2A. The authors declare no competing interests.

